# Treatment-mediated selection of lethal prostate cancer clones defined by copy number architectures

**DOI:** 10.1101/2022.09.01.506183

**Authors:** A M Mahedi Hasan, Paolo Cremaschi, Daniel Wetterskog, Anuradha Jayaram, Stephen Q Wong, Scott Williams, Anupama Pasam, Anna Trigos, Blanca Trujillo, Emily Grist, Stefanie Friedrich, Osvaldas Vainauskas, Marina Parry, Mazlina Ismail, Wout Devlies, Anna Wingate, Stefano Lise, Shahneen Sandhu, Gerhardt Attard

## Abstract

Despite initial responses to hormone treatment, metastatic prostate cancer invariably evolves to a lethal state. To characterize the intra-patient relationships of metastases that evade treatment, we performed genomewide copy number profiling and bespoke approaches targeting the androgen receptor (AR) on 142 metastatic regions from 10 organs harvested post-mortem from nine men who died from prostate cancer. We identified diverse and patient-unique alterations clustering around the *AR* in metastases from every patient with evidence of independent acquisition of related genomic changes within an individual and, in some patients, the co-existence of *AR*-neutral clones. Using the genomic boundaries of pan-autosome copy number change, we confirmed a common clone of origin across metastases and diagnostic biopsies; and identified in individual patients, clusters of metastases occupied by dominant clones with diverged autosomal copy number alterations. Autosome-defined clusters were characterized by cluster-specific *AR* gene architectures that in two index cases were topologically more congruent than by chance (*p*-values 0.03, 3.07×10^-8^). Integration with anatomical site suggested patterns of spread and points of genomic divergence. Copy number boundaries identified treatment-selected clones with putatively distinct lethal trajectories.

**Statement of significance:** Lethal prostate cancer evolves from a single clone of origin and upon a treatment-mediated selection, progresses to lethal disease via a limited number of related clones harboring patient-unique androgen receptor gene architectures.

## Introduction

Deciphering the conundrum of treatment resistance in metastatic epithelial malignancies is a major unmet medical need. Prostate cancer shows high response rates to androgen deprivation therapy (ADT), but relapses often occur after an average of two years (1). Selection of clones harboring *AR* amplification or mutations (2, 3), structural rearrangements, splice variants and a plethora of events consistent with treatment-mediated selection to maintain AR activity despite medical efforts to inhibit it result in an often rapidly lethal state that remains poorly understood (1). AR alterations in liquid or tissue biopsies associate with shorter responses to second-line next-generation hormonal treatments (4, 5) and new drugs are in development to target aberrant AR (6–8). However, heterogeneity of AR alterations across metastases could create a challenge that complicates their utility as a biomarker or therapeutic target. We therefore aimed to characterize *AR* genomic complexity across spatially-separated lethal metastases and then by using pan-autosome copy number features, sought to evaluate the relationships of intra-patient metastases to inform on their evolutionary and metastatic trajectories.

## Materials and Methods

### Patients and rapid warm post-mortem program

All the patients (N=9, median age, 68, all male) involved in this study agreed to participate in a communitybased rapid autopsy (Cancer Tissue Collection After Death, CASCADE) program described previously (9). The inclusion criterion for this analysis was that the patient died from metastatic castration-resistant prostate cancer (mCRPC). All cases analysed are included in this report. The number of samples to be included in the analysis was not pre-defined based on power calculations and we aimed to include the maximum number with sufficient tumor content. The CASCADE program was sponsored and conducted by the Peter MacCallum Cancer Centre, Melbourne, Victoria, Australia and approved by the Human Research Ethics Committee of the Peter MacCallum Cancer Centre, Melbourne (HREC approval numbers: CASCADE 13/122). Tumor samples were obtained post-mortem, and site of biopsy was carefully annotated and photographed. Where possible, formalin-fixed paraffin-embedded (FFPE) tumor samples acquired whilst the patient was alive were retrieved. There was no randomization or blinding involved. The experimental study design is summarized in **Figure S1**. All patients provided informed consent.

### Determining *AR* gene copy number

The ACE and QDNASeq R package (10, 11) was used to determine the autosomal copy number profiles from the low coverage WGS BAM files using a bin size of 500 kb (details in the supplementary methods). We used the cellularity and ploidy values from our in-house approach (details in the supplementary methods) for this calculation, where possible. Validation of *AR* copy number was performed using droplet digital PCR as described previously (12) (**Supplemental Methods, Figure S2**).

### Somatic mutation calling and annotation

Somatic mutations were determined from whole genome, whole exome and targeted deep sequencing reads using GATK4 Mutect2 (13) with default parameters. A panel of normal, beside matched normal samples in each patient, from the Broad Institute (gs://gatk-best-practices/somatic-b37/Mutect2-WGS-panel-b37.vcf) was also used to filter out the false positives and gnomAD vcf file (https://gnomad.broadinstitute.org/downloads), also from the Broad Institute, was used to further filter the mutation calling based on population allele frequencies of common and rare alleles. Then the somatic mutations were further filtered using FilterMutectCalls function as per the GATK best practices. Somatic mutations were then annotated using dNdScv (14) and the synonymous variants were filtered out in the subsequent analyses.

### Somatic structural variant calling

Delly v-0.7.8 (15) was used to determine somatic structural variations in the metastatic samples against the matched normal sample for each patient using high coverage WGS or targeted sequencing data (details in the supplementary methods). Cancer cell fraction (CCF) was calculated for targeted sequencing derived structural variants applying a previously published method (16).

### Determining relationship of metastases

Consecutive genomic bins with similar normalized reads counts derived from copy number analysis were merged to form copy number segments. A revised copy number profile of each patient was determined by the copy number value of each segment at its boundary (henceforth termed as “Transition points”). We employed SCRATCH (Start of Copy number change for Relationship Assessment and Testing Clone Histories) method to determine the relationship of tumors in a patient (details in the supplementary methods). “Silhouette” method, as implemented in the R package “cluster” (17), was used to identify the optimal number of clusters in each SCRATCH relational network. A cluster having less than two samples, was merged with the adjacent cluster sharing the most recent common ancestor on the SCRATCH relational network.

### Clonal decomposition and validation of SCRATCH

Clonal decomposition was performed on each metastatic sample using Sclust (18) (details in the supplementary methods). Somatic mutations that belong to cluster 0 in <sample>_mclusters.txt output files were taken as the dominant population in each metastatic sample and further annotated for the impact of mutations using dNdScv (14). Non-silent mutations were considered when an intersection of such mutations between a pair of metastatic samples from a patient were calculated. This intersection value was further normalized by the smaller total number of the non-silent clonal mutations belonging to a sample between the pair of comparisons.

### Congruence analysis

Baker’s Gamma correlation coefficient (19) was used as a measure of association (similarity) between two trees of hierarchical clustering (dendrograms). From the R package “dendextend”, the function cor_bakers_gamma() was used to calculate this correlation coefficient. For calculating the Congruence Index (*I_cong_*) (20), precompiled scripts were used upon receiving from the author through a personal communication. A *p*-value of 0.05 was taken as the cutoff for determining if two trees are more congruent than by chance.

### AR score

Gene-wise count data were filtered for a minimum of 10 reads in at least 90% of the samples. Then the count data was quantile normalized using voom() function of Bioconductor package “Limma” (21) and z-sores were calculated for the *AR*-response genes (N=20) described by Hieronymus and collaborators (22). We assigned a ternary scoring system following the algorithm below –

AR score for each of the genes up-upregulated by AR (22):

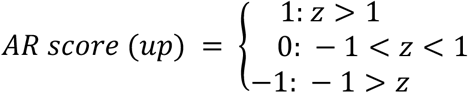

On the other hand, for genes reported as downregulated, a converse scoring algorithm was implied. Finally, all the AR scores (N=20) were aggregated to calculate the final AR score.

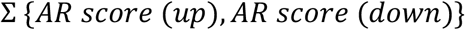

### Statistical analysis

All quantification and statistical computing (done using R software, unless otherwise stated) were performed as described in the figure legends. In brief, median coverage of the WGS samples was determined using Picard (https://broadinstitute.github.io/picard/). The RNA-seq count was quantile normalized using “voom” function in R/Bioconductor package “Limma” (23) while calculating AR scores (to determine *AR* gene downstream transcriptional activity). To compare AR downstream transcriptional activities between tumors with *AR* copy number gain or normal, Kruskal-Wallis test was performed. Wilcoxon test was performed to compare the node distances on SCRATCH relational network between clusters or organ types. In addition, Silhouette (24) analysis was used to study the separation distance between the SCRATCH-defined clusters. Congruence analysis between autosomal- and chromosome X-based SCRATCH relational networks relationships was performed using both Baker’s Gamma index (19) and Congruence Index (*Icong*) (20), using a *p*-value of 0.05 as the cutoff.

### Data and code availability

The whole genome and RNA sequencing data generated as part of this study will be uploaded to the European Genome-Phenome Archive and will be available to the readers following review by the CASCADE data access committee.

## Results

### High selective pressure for structural alterations involving the *AR* gene region

We performed rapid post-mortems on nine men who died from mCRPC. Over the course of their treatment, between 2001 and 2017, all men had developed treatment resistance, defined by a rise in serum prostate specific antigen (PSA) on second-generation AR signaling inhibitors (abiraterone or enzalutamide, **Figure 1a**, **Table S1**). In total we harvested 182 fresh frozen tumor samples from 10 different organs. We also collected plasma from eight men at death and retrieved 26 archived formalin-fixed paraffin embedded (FFPE) tumor samples from seven of the men. These FFPE tumor samples were obtained at diagnosis or after development of castration-resistance, either for palliation or as a requirement for participating in a clinical trial.

**Figure 1.**
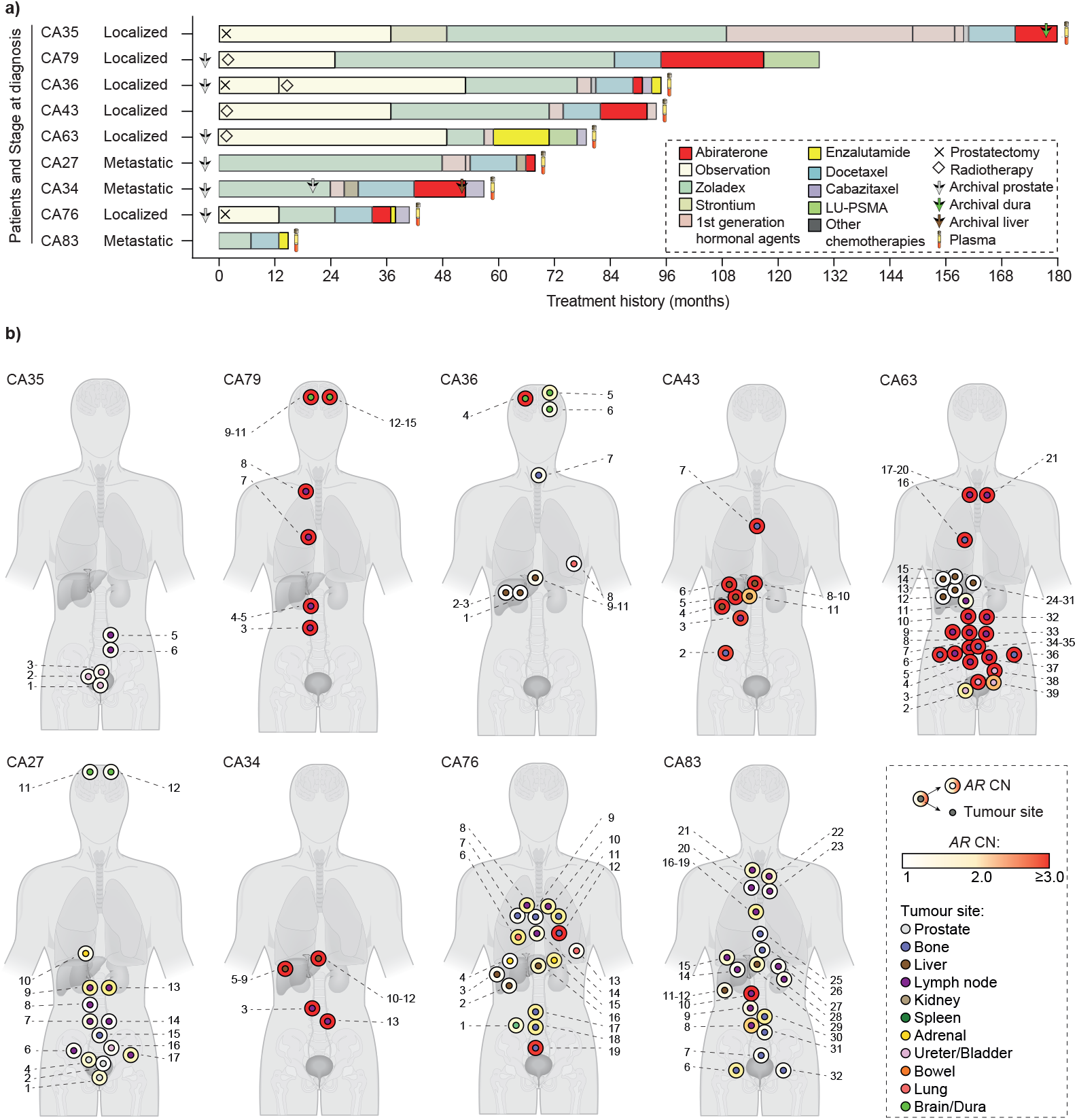
Treatment regimen, anatomical positions and AR copy number status of metastases harvested post-mortem. a) Horizontal stacked bars denote the treatment regimen administered, color codes in the legend. Patients are arranged based on the time from diagnosis to death. Arrows denote the time of tissue biopsy (prostate, brain or liver) at initial diagnosis or as part of standard-of-care. Plasma was collected post-mortem from eight patients, denoted by blood collection tubes. b) AR copy number state (outer circle) and approximate anatomical positions of tumors with tumor content >=0.2.

Copy number alterations are common in advanced prostate cancer (25). We first used shallow whole genome sequencing (median coverage: 0.37X, range: 0.07X-5.8X, **Table S2**) to estimate tumor fraction and obtain copy number profiles (see **Figure S1** for experimental study design). For tumor fraction estimation, we used the consensus of two approaches: a previously published method implemented in the ACE package (11) and, secondly, a bespoke strategy that derived a measure of tumor content based on the allelic imbalance at heterozygous single nucleotide polymorphisms (SNPs). To maximize the accuracy of the latter approach, we concurrently performed whole genome sequencing (median coverage: 23.5X, range: 14.14X-32.62X) on the nine patients’ germline samples (described in Methods). This led us to exclude samples with a tumor fraction of less than 0.2 (not including), namely 40 (22%) fresh-frozen post-mortem, three plasma (43%) and nine formalin-fixed archival (35%) samples. Of the resultant formalin-fixed samples, for six patients we had standard-of-care diagnostic prostate biopsies, for two patients we had only CRPC biopsies (brain and prostate) and for one patient we had a diagnostic biopsy, a prostatectomy sample obtained 18 months after the first biopsy sample was taken and a mCRPC liver biopsy six months before death (**Figure 1a**).

We then extracted *AR* copy number from the remaining 142 post-mortem samples. Of these, 59 (42%) had gain of the *AR* (defined as copy number ≥2 and involving <80% of Xq, also see Methods, **Figure 1b**), confirmed by using droplet digital PCR (ddPCR) or targeted NGS (**Figure S2**), including at least one metastasis from seven out of nine men. We observed distinct groups of metastases with overlapping patterns of chromosome X copy number architecture showing intra- and inter-patient diversity (**Figure S3**).

We also noted that variably large areas of chromosome X showed copy number gain (**Figure 2**). Invariably when copy number change occurred, it involved the *AR* and in metastases from CA34, CA43, CA63 and CA79, also its associated (centromeric) enhancer, which in its gained state, was recently shown to be associated with resistance to abiraterone or enzalutamide (26, 27).

**Figure 2:**
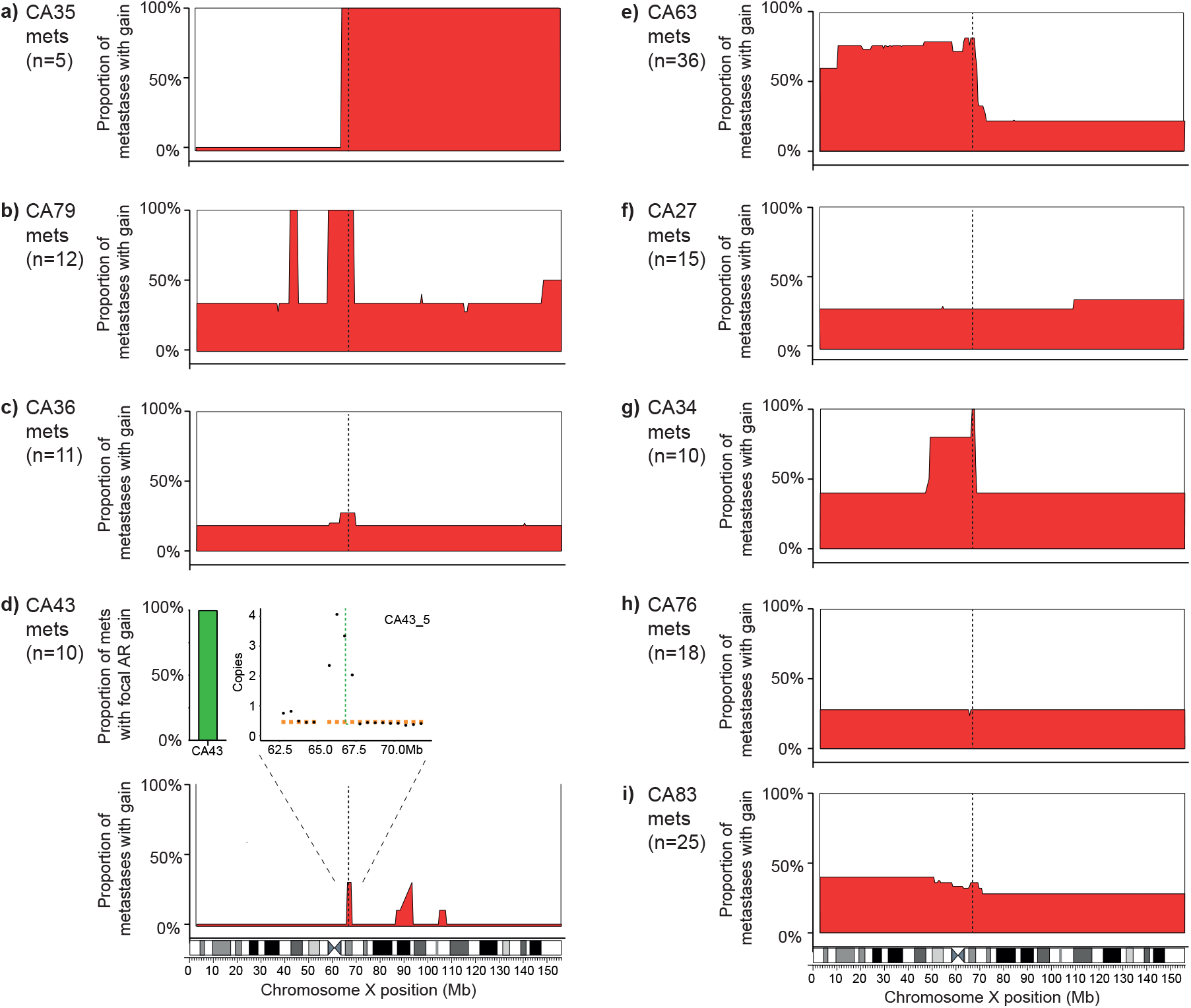
Chromosome X architecture captures intra- and inter-patient diversity. a-j) Skyline plots showing the proportion of samples in each patient with copy number gains (>=2 copies, along y-axis) at individual bins (500kb wide) across chromosome X (along x-axis) (**Table S6**). Number of metastatic samples per patient (n) is provided for each panel. Vertical dotted lines denotes position of AR gene and associated enhancer region. (i) Inset used to show focal gain in AR, not evident at low resolution in the skyline plot. Orange squares depict mean copy number for contributing segment(s) and black dots represent copy number of each bin.

We then used the above information to select two to six metastases from each patient for resequencing at a higher depth of coverage (total: 22; median coverage: ~56X; range: 27X – 82X, **Table S2**). Focusing on chromosome X in these data, we identified breakpoints that were unique to each patient and occurred within a large genomic area around *AR* (**Figure 3a**). This is of relevance to efforts of utilizing *AR*-associated breakpoints as biomarkers for tracking of tumor clones and treatment selection. We noted overlapping breakpoint occurrences in spatially separated metastases (for example, CA34 right and left liver lobes; and CA63 vertebra and soft tissue of rib). We studied this further on samples from patient CA34, which had the highest density of breakpoints converging on *AR*, by using high-coverage NGS custom probes (**Figure 3b**). The break-point associated with a *TMPRSS2-ERG* fusion was present in all tumors analyzed and was putatively an early event that preceded acquisition of structural rearrangements in the proximity of *AR*. The clone most abundant in liver metastases and its “gatekeeper” lymph node harbored at least six break-points unique to this patient around *AR* (BP1 to BP6), admixed at varying proportions. We also identified evidence of subclonal seeding of two external lymph node metastases (CA34_3, CA34_2) and a prostate biopsy (CA34_1) with varying proportions of these six break-points. The sum of the abundance of each break-point at these later tumor sites was less than the total cancer cell fraction, suggesting they occurred together with genomically-distinct sub-clones that did not harbor rearrangements involving the *AR* region.

**Figure 3:**
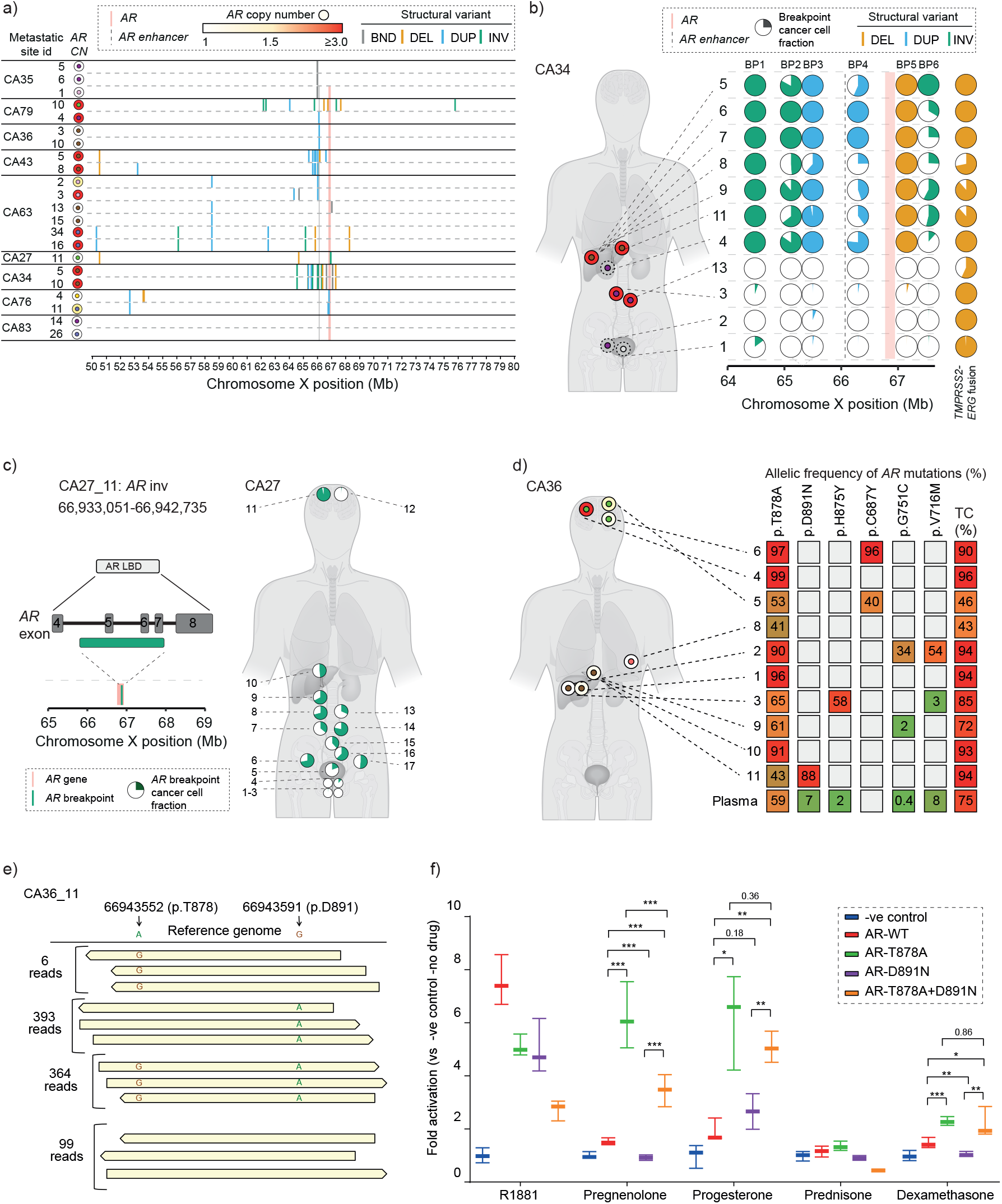
High selective pressure for genomic alterations involving the AR gene region. **a**) Breakpoints at the start of structural variants (BND: translocation, DEL: deletion, DUP: duplication/gain and INV: inversion) detected in 22 high-coverage (~60X) whole-genome next-generation sequencing (NGS) on samples from ten individuals, showing a 30 Mega-base region of chromosome X around AR gene (**Table S7**). **b**) <u>Left panel</u>: anatomical positions of metastatic samples (inner circle) and their AR copy number (outer circle) are shown for CA34 (**Table S7**). Dashed outer circles depict samples with tumor content <0.2. <u>Right panel</u>: pie charts depicting the clonality (by cancer cell fraction, CCF) of breakpoints in proximity of the AR gene and on chromosome 21 at position of TMPRSS2:ERG fusion detected by high-coverage targeted sequencing on 11 samples. AR and associated enhancer are depicted with vertical salmon and blue solid lines, respectively. **c**) <u>Left panel</u>: High-coverage custom NGS confirmed a copy number neutral breakpoint (for an inversion event) in exon 3 of AR gene in patient CA27 which resulted in a ligand-independent, constitutively activated AR splice variant. <u>Right panel</u>: the distribution of the sub-clonal breakpoint and associated CCF at different anatomical sites are shown with pie charts (**Table S7**). **d**) Pathological mutations detected in CA36 metastases using high-coverage targeted sequencing. Allelic fractions and tumor content are indicated by color and number, respectively. Anatomical sites of tissue sampled (left panel, color coded inner circles as for Figure 1b) are shown with AR copy numbers (outer circles). **e**) Independent acquisition of AR mutations in a liver metastasis (CA36_11) detected using amplicon-based, high-coverage targeted sequencing. The supporting reads confirming each allele type shown on left: wild-type alleles are shown on top and mutated alleles are shown along the cartoon reads. **f**) Reporter-luciferase assay showing activation of wild-type and mutant AR (T878A and D891N, individually and combined) by clinically relevant ligands (R1881: Synthetic Androgen). Significance tested with non-parametric t-test; significance code: ‘***’ <=0.001, ‘**’ <= 0.01, * <=0.05).

There was no evidence of *AR* copy number gain in CA27 metastases. However, we detected an inversion involving exon 5 to 7 of *AR* that was previously shown to result in an *AR* splice variant that lacked ligand-binding capability and was constitutively activated independent of ligand (**Figure 3c**) (28). Using customized high-coverage targeted NGS and ddPCR, we confirmed the copy-number neutral breakpoint to be present in every metastasis and in one of the four prostate tumor regions at varying fractions (at sub-clonal level) admixed with *AR* neutral cells (**Figure 3c**). Overall, we effectively confirmed diverse and patient-unique alterations clustering around the *AR* in metastases from every patient, supporting the potent selective pressures for *AR* aberrant clones in men receiving hormonal therapies.

### Convergence of independent *AR* somatic point mutations

Hormone treatment pressures can also select clones harboring *AR* somatic point mutations (3). We performed ~1400X targeted NGS designed to capture *AR* coding sequences and recurrent prostate cancer mutations and in total detected seven unique non-synonymous functionally-active mutations within *AR* coding regions. Of these, six (encoding for T878A, C687Y, H875Y, D891N, G751C, and V716M) occurred in CA36 and one (E710G) in CA43. In CA36, pT878A was detected in every metastasis (**Figure 3d**). Prior studies suggest this mutation is the most common in mCRPC patients, detected in 10-15% of the cases (28, 29). We detected all *AR* somatic mutations in plasma that were harbored in extra-cranial metastases but not pC687Y that was detected solely in the brain metastases.

We hypothesized that all metastases could either have been seeded by the same clone harboring a T878A mutation or the clones independently acquired it after metastatic spread. In our search for evidence supporting either hypothesis, we identified in one of the liver metastases (CA36_11) alleles with mutation(s) encoding for D891N or T878A or both (**Figure 3e**). The presence of tumor sequencing reads with a wild-type allele at this position (X:66943552) suggests this mutation occurred after metastatic seeding, despite its presence in every metastasis, and emerged independently in the lineages with D891N or without. An alternative explanation could be a mutation reverting to the wild-type but this appears unlikely. To further investigate the biological pressures for selection of these two *AR* mutations, we used a reporter luciferase construct in prostate cancer cells transfected with AR wild-type or a combination of the detected mutations. We found that acquiring a T878A change at a D891N altered allele conferred increased activation by a range of ligands including progesterone and pregnenolone (**Figure 3f**), previously suggested as contributing to resistance against abiraterone (12, 30). We also observed evidence of a second independent event in the right dural metastasis (CA36_4) with an amplification of the T878A *AR* mutant allele (**Figure 3d**). We found no evidence for differential ligand-activation by AR E710G compared to wild-type (data not shown). In CA43, we observed gain of the mutant pE710G allele in four of six and of the wild-type allele in two of six metastatic samples (**Table S3**).

### Autosomal copy number transition points define tumor relationships

To interrogate the evolutionary paths followed by groups of metastases with different *AR* gene architectures, we then deliberately focused on copy number change that occurred in the autosome. In several cancers, parts of the genome are affected by copy number alterations that can show an evolutionary order to their occurrence across patients and cancer types (31). In prostate cancer copy number change is common (32) and, although involved regions are repeated across cancers, the starting event that leads to each change could be different in every patient’s tumor. We posited that the junction in the genome where copy number change occurred could therefore offer an opportunity for “tumor fingerprinting” to track relationships over time and space. Using copy number profiles derived from whole-genome sequencing, we identified the boundaries at which a change in the copy number of adjacent autosome segments occurred (henceforth referred to as “transition points”).

We investigated the co-occurrence of transition points in temporally-separated archival formalin-fixed biopsies with tumors harvested post-mortem. Whilst we observed sharing of individual transition points across metastases from different individuals (**Figure S4**), we uniquely observed that a great proportion of transition points in archival samples from an individual were shared with all metastases harvested postmortem from the same patient (putatively patient-common or truncal transition points) (**Figure 4a and S4**). This reaffirms a common clone of origin in metastases at death (33) and now shown here, the same clone of origin in diagnostic biopsies.

**Figure 4.**
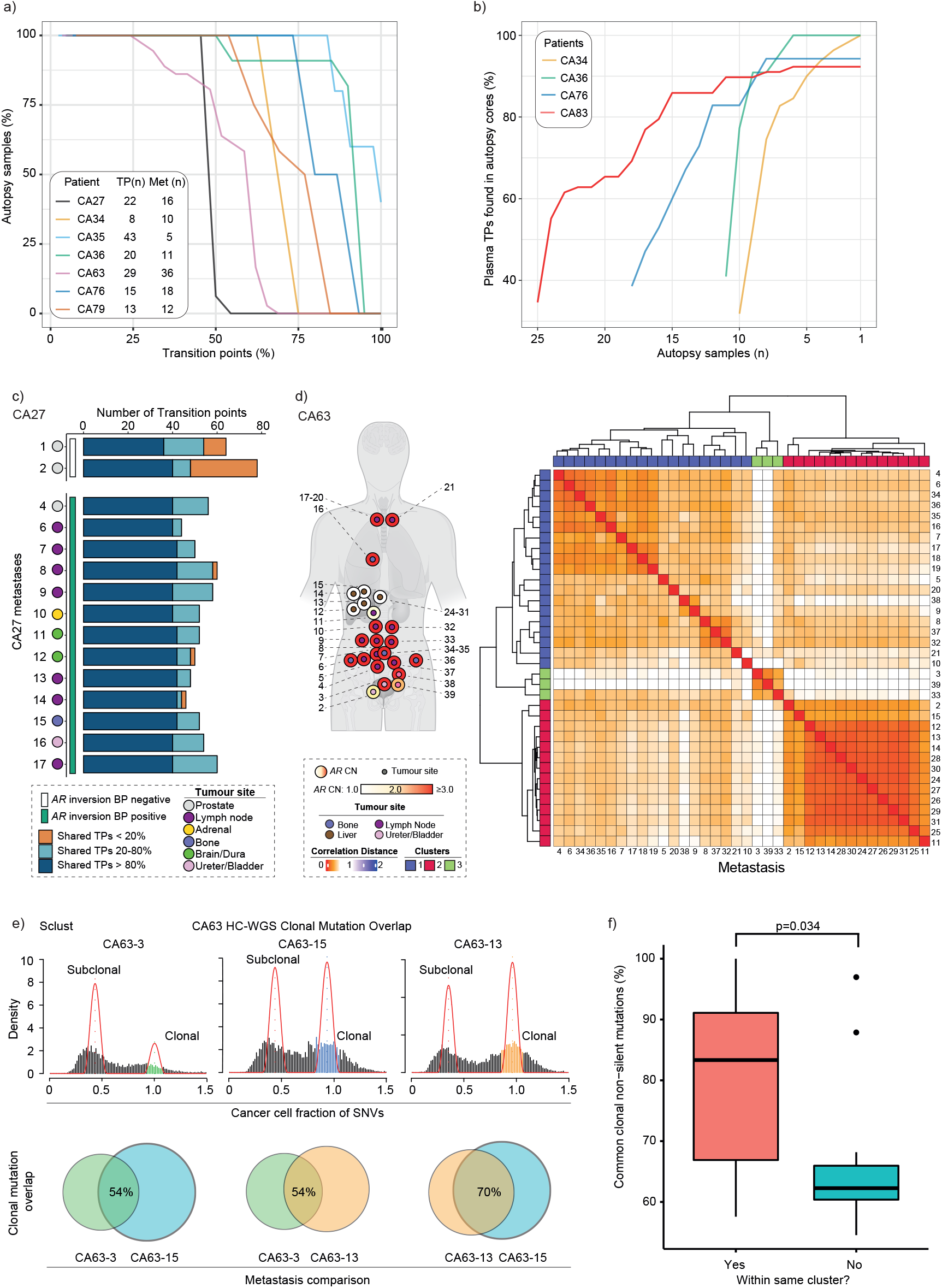
Copy number transition points confirm the same origin of lethal prostate cancer and diagnostic biopsies and define the relationships of lethal metastases. **a**) Line plots showing the percentage of autopsy samples harboring transition points detected at diagnosis from the same patient (n=7) against the respective set of transition points (as percentage). **b**) Proportion of transition points detected in plasma (tumor content >= 0.2), collected post-mortem, are plotted as a function of the number of post-mortem metastatic samples from the same patient (n=5) (**Table S8**). **c**) Stacked bars show different percentages (<20%, 20 to 80% and >80%) of shared copy number transition points among autopsy samples in CA27 (**Table S8**). Two prostate tumors (1 & 2) do not share a pathogenic AR inversion break point and display a higher tumor-unique transition points, while the remaining prostate tumor (CA27_4) shows more homogeneity of shared transition points with other distal metastases and, in unison, share the break point. **d**) <u>Left panel</u>: Metastatic samples harvested postmortem in CA63 are depicted with anatomical position (inner circle) and AR copy number (outer circle). <u>Right panel</u>: Post-mortem metastatic samples form three distinct clusters by applying the SCRATCH clustering algorithm (described in methods). The color scheme of the heatmap is based on the correlation distances calculated using copy numbers at transition points. **e**) Illustrative figure showing how clonal mutations (**Table S5**), detected using Sclust, form a distinct peak at cancer cell fraction of 1.0. Intersection of non-silent clonal mutations between two samples were chosen and normalized by the total number of the smaller set of such mutations (described in methods). **f**) Box plot showing the distribution of percent common clonal non-silent mutations by assignment of post-mortem samples in a cluster. Willcoxon non-parametric test was used to measure significance of difference between those two distinct groups, as shown.

When we compared plasma at death and metastases obtained post-mortem for four patients (CA34, CA36, CA76 and CA83), to our surprise, we identified that in all cases >90% of transition points in plasma were detected in the metastases analyzed, suggesting that metastatic sampling captured the majority of DNA shedding clones (**Figure 4b**). We then compared transition points in CA27 metastases and the prostate tumor that were intra-*AR* break-point positive with the two prostate samples harvested post-mortem that were breakpoint negative. We identified clonal diversity between these two groups where although all tumors shared the majority of transition points, the two break-point negative prostate tumors had a high proportion of transition points that were unique to their respective clones (**Figure 4c**).

In addition to truncal transition points shared across metastases from the same patient, in some patients we noted that transition points were shared across distinct groups (or clusters) of metastases. We therefore developed a hierarchical clustering-based algorithm (we refer to as Start of Copy number change for Relationship Assessment and Testing Clone Histories, SCRATCH) to determine the relationship of metastases (hence named “SCRATCH relational network”) based on the correlation of the copy number values at transition points (**Figure 4d, Table S4**). This framework attempted to split metastases into groups that maximized the inter-cluster and minimized the intra-cluster distances based on the presence of copy number differences (see Methods). We focused on the patient with the most metastases and assumed each cluster had a distinct dominant clone. We found evidence of three clusters of metastases occupied by dominant clones in CA63 (**Figure 4d**). We also noted that in three cases with plasma collected postmortem (out of four), the SCRATCH relationship network assigned plasma as having a great similarity to liver metastases (**Figure S5**).

### Dominant autosome copy number changes converge with clonal mutations

In a patient, if metastases in the same SCRATCH-defined cluster had evolved separately from those in a different cluster, we expected that they would share the same clonal mutations. To test this, we first confirmed metastatic samples were assigned to the same cluster regardless of sequencing depth (**Figure S6**). We then hypothesized that the correlation distance (or node distance) between a pair of samples based on transition points would correlate with the overlap in non-silent clonal mutations arising from the dominant clone. We extracted the node distance for metastases with high-coverage WGS and using a previously described tool, Sclust (18), that controls for tumor fraction and ploidy, defined clonal mutations in each patient (**Table S5**). We then calculated the intersection of total non-silent clonal mutations in pairs of metastases (N=24) from the same patient, controlled for total number of the smaller set of mutations between the compared metastases, and confirmed a difference in the number of clonal mutations in metastases in the same SCRATCH-defined cluster from those in other clusters (*p*-value 0.034) (**Figure 4e & f**). This suggests that deriving tumor clone relationships using copy number transition points is consistent with assumptions made using clonal mutation analysis.

### *AR* structural alterations occur in a limited number of established clones with distinct autosome copy number profiles

Given *AR* alterations are selected by hormone treatment, we posited that these are relatively late events in prostate cancer evolution, emerging from established clones at the development of resistance to ADT. To investigate this, we used congruence models to test whether a cluster of metastases derived from autosome transition points was more likely to have the same *AR* structural alterations. By separately using both the Baker’s Gamma index (19) and Congruence Index (*I_cong_*) (20) on patient CA34 and CA63, we observed that clusters of metastases defined by their autosome indeed associated with metastases in the same cluster defined by their chromosome X copy number profile, irrespective of anatomic site (**Figure 5a & b**). These relationships were topologically more congruent than by chance (*p*-value 0.03 and 3.07×10^-8^ for CA34 and CA63, respectively). This was also confirmed by informal annotation that metastases in the same cluster had the same *AR* copy number status.

**Figure 5:**
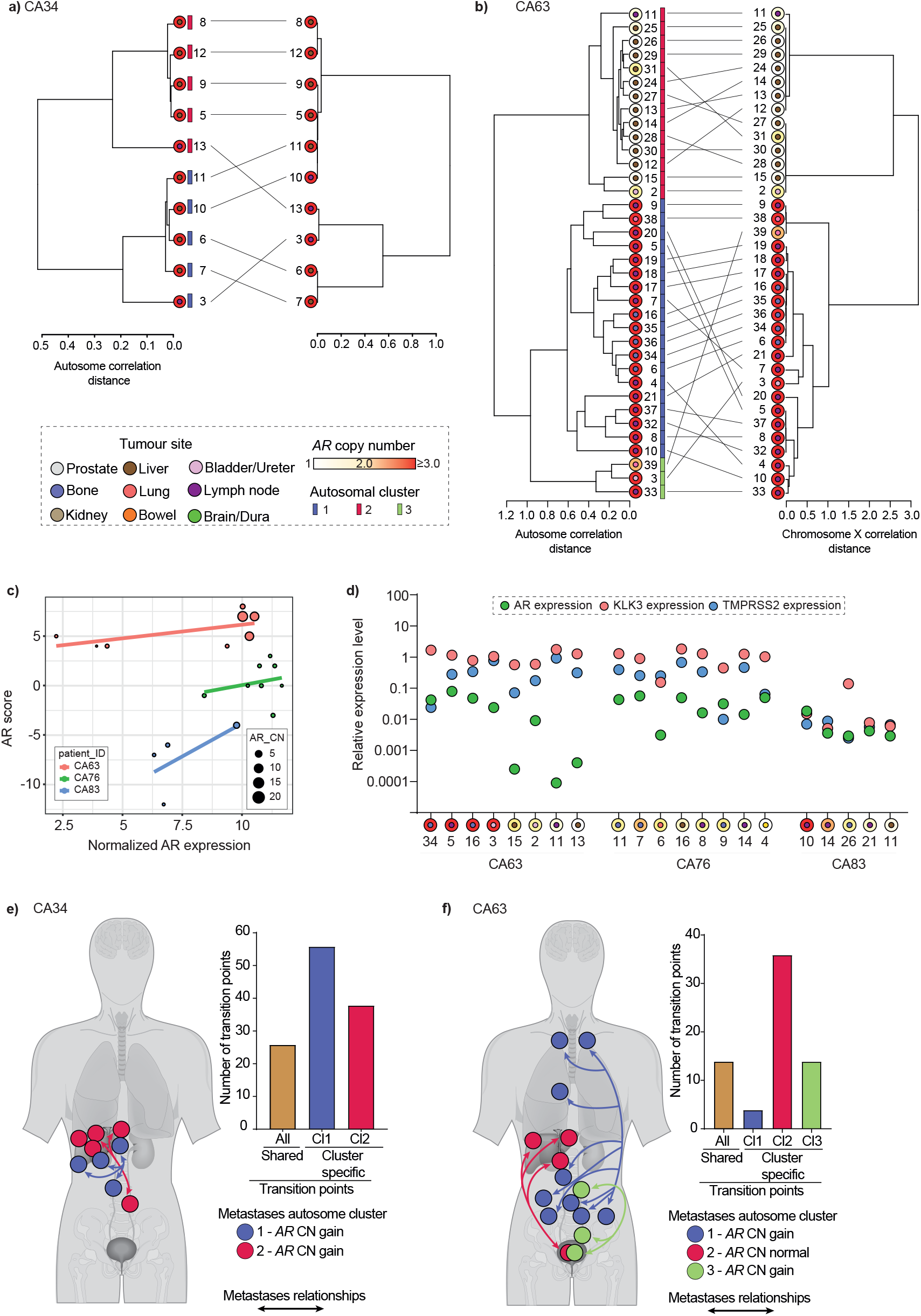
Congruence of autosomal copy number and chromosome X derived relationships suggests selection of AR alterations in established clones. Tanglegrams for (**a**) CA34 and (**b**) CA63 with autosomal SCRATCH relationship network of metastases on the left and chromosome X-based SCRATCH relationship network on the right. Grey lines between relational networks are connecting the same metastatic samples between two SCRATCH determined metastatic relationship. (**c**) AR scores plotted as a function of AR expression (‘voom’ normalized, x-axis) for CA63, CA76 and CA83 (**Table S9**). Linear regression lines are drawn for each patient. (**d**) Expression of AR and it’s regulated genes KLK3 and TMPRSS2 detected by ddPCR, are plotted on a logarithmic scale (y-axis) for three patients (CA63, CA76 and CA83) (**Table S9**). AR copy number and organ sites are depicted in nested circles as depicted in Figure 1b. **e & f**) Cartoons of postulated metastatic evolutionary relationships of dominant clones for CA34 (two clusters) and CA63 (three clusters) are shown with bidirectional trajectories (as reverse migration of metastatic clone(s) cannot be ruled out). Bar charts show common or cluster-specific shared copy number transition points for respective patients.

### Metastatic trajectories to lethal disease

We then integrated AR expression data from RNA sequencing on metastases showing intra-patient differences in *AR* status (N=20 from CA63, CA76 and CA83) based on *AR* architectures and autosome copy number. We also interrogated AR transcriptional activity using a previously-described z-score based method exploiting the expression levels of AR downstream genes (22). There was an overall weak but positive correlation between AR expression and AR score (**Figure 5c**). Moreover, we found no difference in the expression of AR-regulated genes between metastases with and without *AR* copy gain (**Figure S7**). There was a notable difference in AR expression in gained compared to non-gained metastases with the exception of the non-gained bladder tumor (CA63_2, adjacent to an area with gain suggesting the possibility of admixture of *AR* gained and non-gained clones) (**Figure 5d**).

In CA34, two adjacent lymph node biopsies (CA34_3 and CA34_13) clustered with two different sets of liver biopsies showing two dominant clones at death, where each clone contained a considerable number of cluster-unique transition points (**Figure 5e**). Similarly in CA63, of the two biopsies from the same bladder tumor, one (CA63_3) was *AR* amplified and the other was not (CA63_2). Clustering based on their autosome copy number transition points assigned the two adjacent tumors to separate clusters with *AR* copy number matching the respective bladder tumor samples. This suggested that the CA63_3 clone harbored a genomic event that amplified *AR* and at death was the most abundant clone in the bone and lymph node metastases whilst the liver metastases and porta hepatic lymph nodes were populated by the non-amplified clone shared with CA63_2 (**Figure 5f**). The anatomical proximity of the bladder to the prostate and the retained admixture of the two distinct clones in opposite sides of the same tumor lesion could suggest that at least two of the three dominant clones at death in anatomically-distinct metastases originated independently from this site. The patient underwent a period of 48 months observation after surgery, prior to start of ADT. We detected a large number of transition points shared in either bladder area, and all metastases in the same respective cluster suggesting continued copy number evolution prior to spread that we posit occurred in minimal residual disease in the bladder wall that remained after surgery.

## Discussion

We here generated extensive whole genome copy number profiling on an average of 12 metastases and primary tumor samples from nine men who died from prostate cancer after receiving abiraterone or enzalutamide. These molecular data will serve as a resource for the community. We focused on characterizing how AR alterations evolve and co-exist in individual patients. We used copy number transition points rather than direct copy number comparisons as the former can be considered the biological event that gives rise to the latter and could contain sufficient evolutionary information for accurate tracking of metastases. This strategy proved effective on FFPE and plasma DNA samples, that introduces the opportunity to expand in future to studies with multiple temporally-separated samples.

We made a number of unique observations. Firstly, we identified a limited number of two or three dominant clones present at death characterized by distinct *AR* gene architectures, with at least in some cases, coexistence of both *AR* altered and gene neutral clones. This was very notably the case in CA63 with clear anatomical delineation of *AR*-gained and *AR*-neutral metastases. The *AR*-neutral metastases showed similar AR activity scores compared to gained metastases. We also noted that there was no evidence of a predisposition of specific anatomical sites (for example the liver) to a specific *AR* status: for example, we observed *AR*-altered or *AR*-neutral hepatic metastases. Secondly, despite this intra-patient heterogeneity, *AR* alterations in clusters of metastases from the same patient were commonly in the same genomic alteration type. For example in CA36, despite evidence of independent acquisition, all metastases harbored one or more functionally-relevant *AR* mutation; in CA27, all metastases analyzed harbored sub-clones with the same *AR* structural variant. Similarly, CA34, CA79 and CA43 clusters had at least two different patient-unique *AR* genomic architectures achieving similar levels of *AR* copy number gain, and in CA34, we identified different frequencies of distinct break-points suggesting multiple sub-clones that had independently acquired structural change associated with similar levels of *AR* amplification. This could justify classification of patients by *AR* gene class for therapeutic intervention, as for example is being attempted for patients harboring *AR* mutations (7, 8). Overall, whilst we identified greater intra-patient diversity than described previously, the uniformity across clusters of metastases dominated by a single clone contextualizes these previous results that might have primarily obtained samples from the same cluster (34). Thirdly, we used autosome copy number transition points to define the relationships of metastases and confirmed a common clone of origin in lethal metastases and the original diagnostic biopsy. Similar approaches have been described previously (35, 36) but our report describes a scalable analysis pipeline that can be implemented in whole-genome sequencing data of variable coverage and from samples of variable quality. Moreover, the patient-unique transition points common to both biopsy and autopsy sample types could be leveraged for tracking of metastatic clones throughout the patient’s duration of disease. Fourthly, using this framework, we identified a high congruence between autosomal copy number acquisition and chromosome X copy number changes, mostly clustered around the *AR*. We hypothesize that the latter are selected at institution of ADT that exerts a strong treatment-mediated selection pressure and serves as an evolutionary node through which a limited number of clones emerge, characterized by enrichment for (but not exclusively) *AR* gene structural alterations.

Our study also has some limitations. Transition points extracted from our data are biased towards events occurring at a high abundance in a tumor sample and therefore focus primarily on the “dominant” or most abundant clone. We hypothesize this clone has fitness advantages that confer resistance and we are currently unable to comment on less abundant sub-clones. Despite a large number of metastases harvested at post-mortem, we have included relatively few temporally separated samples, so evolutionary inferences are unable to confirm “when specific clones were selected”. For example, in CA27, we are unable to ascertain whether the resistant *AR*-altered clone emerged in the prostate and then seeded distant metastases or spread to the prostate after start of treatment. The former scenario would support the use of local treatment of the prostate, recently shown to improve long-term survival in low-volume, metastatic patients (37). Similarly, although in CA63, it appears more likely that the resistant dominant clones diverged and spread from the bladder, it is also possible that one or all independently spread to the bladder. In the absence of multiple temporally-separated sequential samples, these patterns may be challenging to distinguish. Also, a few transition points unique to the archival samples could have escaped detection in metastatic samples due to additional occurrences of structural events, later on, which could mask the common transition points while segmenting the genome with bins of similar copy number. Nonetheless, those later events in metastases could indicate the divergence of metastases leading to treatment resistance. The current size of the cohort limits testing of associations between patterns of copy number clustering at death and metastatic status at presentation or distinct treatment sequences. The collection of tumors at death is pragmatic and dependent on feasibility and presence of visible tumors. We, therefore, might have over-represented clones in more accessible regions such as the liver. Moreover, as the very definition of a transition point is a function of the bin size that is selected during copy number analysis and application of segmentation algorithms, we have smoothened the data and could have grouped metastases with different breakpoints under a single copy number segment that eventually resulted in copy number changes with apparently the same transition points. Given the resolution of these data, there could be misassignment of the terminal nodes of our hierarchical clustering. Nonetheless, the dominant nodes showed a high congruence with orthogonally obtained detailed *AR* analysis. Finally, by assessing the most abundant copy number changes in a whole biopsy, our study focuses on the common clone in an individual metastasis and does not exclude the possibility of admixture of less abundant sub-clones between different organs (38, 39).

In conclusion, we identified that a limited number of established clones with common copy number transition events are selected by treatment and are characterized by unique *AR* gene architectures. Future studies integrating temporally separated and multi-dimensional data could more closely inform on when this clonal separation occurred and in doing so identify therapeutic opportunities. Overall, our study provides novel insights on the genomic evolution of prostate cancer to a lethal, drug-resistant phenotype. We hypothesize that whilst uniformity of selected *AR* alteration class supports a dependence in an individual on specific resistance escape routes, which could be disrupted by appropriately-timed targeted intervention, the co-existence of *AR*-neutral metastases suggests established clones without *AR* gene disruption can concurrently survive androgen deprivation and progress to form lethal metastases.

## Acknowledgements

We thank all the donors and their families who participated in the Cancer Tissue Collection After Death (CASCADE) program, along with CASCADE investigators, CASCADE Management Committee, all staff at the Victorian Institute of Forensic Medicine, D. Stevens, and Tobin Brothers Funerals.

## Author contributions

Conceptualization: GA, SS

Investigation: GA, SS

Sample acquisition: AP, SS

Methodology: GA, AMMH, DW, PC, SL

Experimentation: DW, AJ

Data analysis: AMMH, PC, SL, DW

Writing manuscript: GA, AMMH, PC, DW, SL

Manuscript review and comments: AMMH, PC, DW, AJ, SQW, SW, AT, SF, OV, MP, AW, SL, SS, GA.

