## Supplementary methods and figures for "Treatment-mediated selection of lethal prostate cancer clones defined by copy number architectures"

### Supplementary Materials

#### Supplementary methods

##### Patients and Rapid Warm post-mortem programs

All the patients (N=9) involved in this study participated in The Cancer Tissue Collection After Death (CASCADE) program. All patients signed informed consent. The CASCADE Program was sponsored and conducted by the Peter MacCallum Cancer Centre, Melbourne, Victoria, Australia. Tumor samples were obtained post-mortem, and site of biopsy was carefully annotated and photographed. The experimental study design is summarized in **Figure S1**.

##### Nucleic acid extraction

Histologically evaluated samples with sufficient tissue material were selected. DNA and RNA from fresh frozen autopsy samples were extracted using the Qiagen Allprep DNA/RNA mini kit (Qiagen) according to the manufacturer's instructions. Archival tumor blocks were retrieved retrospectively, and areas of tumor identified on Haematoxylin and Eosin slides were dissected. Germline DNA was extracted from white blood cells using QIAamp DNA Micro Kit (Qiagen).

##### Digital droplet PCR assay

ddPCR was performed on a QX200 system (Bio-Rad) using the ddPCR Supermix for Probes (Bio-Rad) for copy number analysis and the One-Step RT ddPCR Advance kit (Bio-Rad) for gene expression analysis. Copy number assays were performed for *AR* (Hs04121925_cn, Life technologies), the centromeric chromosome X gene *ZXDB* (Hs02220689_cn, Life Technologies) with *NSUN3* (dHsaCP2506682, Bio-Rad), *HCN1*, *AP3B1* as the reference genes. Gene expression assays were performed for *AR* (dHsaCPE5047114), *KLK3* (dHsaCPE5026548) and *TMPRSS2* (dHsaCPE5051496) using *GAPDH* (dHsaCPE5031597) and *ACTB* (dHsaCPE5190200) as reference genes. PCR reactions were prepared with 2-4 ng DNA or RNA in a total volume of 22 µl and partitioned into ~20,000 droplets per sample with an Automated Droplet generator (Bio-Rad). The PCR reaction was performed and then read on a Bio-Rad QX200 droplet reader using QuantaSoft v1.3.2.0 software for either copy number or gene expression analysis.

**Library preparation and next generation sequencing**

DNA libraries were prepared using the NEBNext® Ultra™ II DNA library prep kit (New England Biolabs) according to the manufacturer's protocol. In brief, 100 ng of DNA input was used for fresh frozen tumor and germline DNA while FFPE DNA inputs ranged from 5-10 ng and included an FFPE DNA repair step (New England Biolabs). All DNA were sonicated on the Covaris E220 (Covaris) to a size of 150-200 bp. Sheared DNA were adaptor ligated, size selected using Agincourt AMPure beads (Beckman Coulter) and PCR amplified using unique dual index primers to reach at least 100 ng of library. Libraries were then pooled and sequenced on a NovaSeq 6000 sequencing system (Illumina) to a desired coverage using 2 x 100 paired-end sequencing for fresh frozen and germline DNA and 2 x 50 paired end sequencing for FFPE DNA. Whole exome libraries were captured using the Roche NimbleGen SeqCap EZ Human Exome Library v3.0 kit and then the library was prepared using KAPA LTP DNA sample preparation kit from Roche, and sequenced at 100 bp paired-end reads on a Illumina HiSeq 2500 sequencing system. Characterization of the *AR* coding region was achieved through targeted next generation sequencing approaches including amplicon enrichment as described previously (*5*) or capture-based enrichment (Integrated DNA technologies). For capture-based enrichment a custom capture probe panel was designed including exonic regions of the genes that were previously reported altered in advanced prostate cancer (**Table S10**) including *AR*. Spike-in probe pools for patient-specific (CA27 and CA34) breakpoints were also designed. 200 ng of whole genome library was used as input for the capture and 10 libraries were pooled into a single capture reaction. Capture reactions were pooled and sequenced on a MiSeq sequencing system (Illumina) aiming for a 100X coverage, using 2 x 75 paired end sequencing. RNA-Seq libraries were prepared using NEBNext® Ultra™ II Directional RNA Library Prep Kit for Illumina as per manufacturer's instruction (with a slight modification to 7 cycles of PCR). Libraries were pooled and sequenced on an Illumina NovaSeq™ 6000 sequencing system (Illumina) to a desired output of 50 million reads per sample.

**Genome sequencing data:**

After basic quality checking using fastqc (https://www.bioinformatics.babraham.ac.uk/projects/fastqc/), reads with adapter contamination were trimmed at the 3' end using skewer (*40*) where the minimum permitted read length was 50 bp. Burrows Wheeler Aligner (BWA) (*41*) was used to map the remaining reads against human reference genome hs37d5 using default options of the bwa mem algorithm. The resulting bam files were coordinate sorted using samtools (*42*) and duplicated reads were removed using picard (https://broadinstitute.github.io/picard/).

**RNA sequencing data:**

RNA-Seq data was mapped using STAR 2.7.9a (*43*) against human reference genome hs37d5 in basic two-pass mode for splice aware read alignment. Count data over gene was generated using HTSeq-count in "union" overlap solution mode utilizing a Gene Transfer Format (GTF) file from Gencode database (https://ftp.ebi.ac.uk/pub/databases/gencode/Gencode_human/release_19/gencode.v19.annotation.gtf.gz) (*44*).

**Somatic mutation calling and annotation**

Somatic mutations were determined from whole genome, whole exome and targeted deep sequencing reads using GATK4 Mutect2 (*32*) with default parameters. A panel of normal, beside matched normal samples in each patient, from the Broad Institute (gs://gatk-best-practices/somatic-b37/Mutect2-WGS-panel-b37.vcf) was used to filter out the false positives and gnomAD vcf file (https://gnomad.broadinstitute.org/downloads), also from the Broad Institute, was used to further filter the mutation calling based on population allele frequencies of common and rare alleles. Then the somatic mutations were further filtered using FilterMutectCalls function as per the GATK best practices. Somatic mutations were then annotated using dNdScv (*35*) and the synonymous variants were filtered out in the subsequent analyses.

**Luciferase reporter assay**

Experiments were conducted as described and referenced previously (*17*). In brief, PC-3 cells were co-transfected with a PSA-ARE3-luc luciferase reporter plasmid and a Renilla luciferase vector plus an empty, *AR*-wildtype or *AR* mutant expression plasmid. Cells were seeded in white opaque 96-well plates and grown in 10% CSS-supplemented phenol red-free RPMI 1640. Cells were then treated with the 0.1nm R1881 or 0.1µM Prednisone, Pregnenolone, Progesterone or Dexamethasone for 16 hours. Luciferase activity was determined using Dual-Glo according to the manufacturer's instructions (Promega) and luminescence was measured on a TopCount plate reader (Perkin-Elmer).

**Somatic structural variant calling**

Delly v-0.7.8 (*31*) was used to determine somatic structural variations in the metastatic samples against the matched normal sample for each patient using high coverage whole genome sequencing or targeted sequencing data. Delly combines short-range and long-range paired-end split reads to call a structural variant (deletion, tandem duplication, inversion and translocation) at a single nucleotide resolution. Structural variants called on both metastatic and matched normal samples were filtered for tumor DNA contamination in the normal sample using a maximum of 0.2 ALT support. Cancer cell fraction (CCF) was calculated for targeted sequencing derived structural variants applying a previously published method (*36*).

**Clonal decomposition and validation of SCRATCH**

Clonal decomposition was performed on each metastatic sample using Sclust (*21*), which calculated the cancer cell fraction (CCF) from the allelic fraction of somatic mutation applying ploidy and local copy number correction. Structural variants were used to add further granularity to the analysis. Somatic mutations that belong to cluster 0 in <sample>_mclusters.txt output files were taken as the dominant population in each metastatic sample and further annotated for the impact of mutations using dNdScv (*35*). Non-silent mutations were considered when an intersection of such mutations between a pair of metastatic samples from a patient were calculated. This intersection value was further normalized by the smaller total number of the non-silent clonal mutations belonging to a sample between the pair of comparisons.

**Tumor content estimation**

In order to determine the tumor content in low coverage samples we exploited the availability of high coverage WGS samples for each patient. The high coverage samples typically had high tumor content and their purity and ploidy status could be characterized with good reliability by combining information from (a) tumor/normal read-depth ratio and (b) phased B-allele frequency at germline heterozygous SNP. Read depth count were calculated genome-wide in bins of 100 kb using ReadCounter() function of HMMCopy v0.1.1 (*45*). Germline samples were genotyped with PLATYPUS v0.8.1.2 (*46*) and phased with BEAGLE5 (*47, 48*) using the 1000 Genomes Project phase 3 reference panel (*49*). Phased B-allele frequency values for tumor samples were calculated in bins of 100K bp, using PLATYPUS v0.8.1.2 (*46*) and in-house scripts. We focused, in particular, on large 'anchor' regions (e.g. chromosome 8p) with low read-depth ratio and with allelic imbalance (e.g. LOH regions). The read-depth levels and B-allele frequency values imposed some constraints on the possible copy numbers of the anchor regions. We determined them by finding a solution that satisfied the constraints across all the anchor regions. Finally, we run the software SEQUENZA v2.1.2 (*50*) and verified that our solution for ploidy, purity and copy number segmentation of the anchor region was among the proposed solutions.

For samples with high tumor purity, in anchor regions with allelic imbalance the phased B-allele values splits into two clearly separated distributions, corresponding to the maternal and paternal chromosomes. We used this to reconstruct the long-range haplotypes in anchor regions and corrected for 'switch-errors' made by the phasing algorithm. We then calculated the haplotype BAF in low coverage samples and determined the tumor content from it. The underlying assumption was that the copy number of the high coverage and low coverage samples was the same in anchor regions. This was often, but not always, the case. We therefore relied on as many anchor regions as possible (minimum of 3) and on a visual inspection for each sample. Finally, we employed the consensus between tumor contents determined by the abovementioned procedure and the ACE estimation for each sample.

**Determining *AR* copy number using ACE:**

QDNAseq/ACE R packages (*11, 33*) were used to determine the copy number profiles from the low coverage WGS BAM files generated from fresh-frozen autopsy samples, plasma at death and biopsy samples using a bin size of 500 kb. To determine the autosomal copy number profiles, we excluded both chromosome Y and mitochondrial DNA and the ploidy was adjusted using median bin segment value, which was the central assumption of ACE. We used a ploidy penalty of 0.5 and lower-cellularity penalty of 0.5 to fit the "squaremodel()" function of ACE as per the author's recommendation. *AR* copy number was determined, as the median copy number of the segment covering the *AR* gene, using getadjustedsegments() and analyzegenomiclocations() functions of ACE. We used the cellularity and ploidy values from our in-house approach (described in the previous section) for this calculation, where possible. We confirmed a high correlation with *AR* copy number estimated using droplet digital PCR (**Figure S2**).

**Correlation assessment of copy number profiles**

During copy number analysis by QDNAseq, consecutive bins with similar normalized reads counts (a.k.a. copy number) were merged to form copy number segments (*33*). A revised copy number profile of each patient was determined by the copy number value of each segment at its boundary (that we termed as “Transition points”). We employed SCRATCH (Start of Copy number change for Relationship Assessment and Testing Clone Histories) method, which implemented hierarchical clustering algorithm on the correlational distances derived from the copy number at the transition points, to determine the relationship of tumors in a patient. The “silhouette” method, as implemented in the R package “cluster” (*34*), was used to identify the optimal number of cluster in each SCRATCH relational network. In the case of a cluster having less than two samples, it was merged with the adjacent cluster sharing the most recent common ancestor on the SCRATCH relational network.

**Supplementary Figures:**

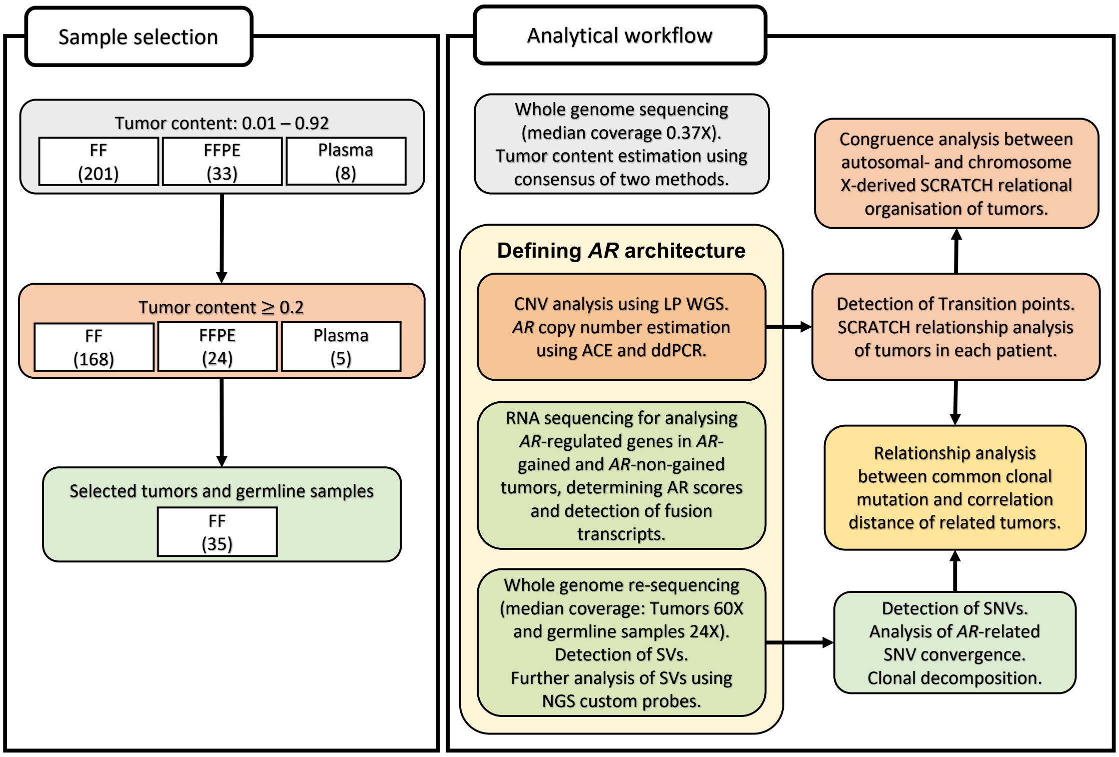


**Figure S1: Experimental Study Design**

**a) Sample summary**: Samples (FF: fresh frozen, FFPE: formalin-fixed paraffin embedded or plasma; sample numbers as shown in parentheses) were subjected to shallow whole genome sequencing (median coverage: 0.37X, range: 0.07X-5.8X). Samples with estimated tumor content ≥0.2 were selected for further analysis, including *AR* copy number determination and genome-wide copy number assessment. Two to six metastases from each patient were chosen as described in the text for re-sequencing at a higher depth of coverage (median coverage: 60X, range 27X-83X). RNA-Seq was performed on 20 samples selected from three patients (CA63, CA76 and CA83).

**b) Analysis summary**: Copy number analysis on low coverage WGS was performed using ACE package, while somatic structural variants surrounding *AR* gene (chrX:50M-80M) were detected from high coverage WGS data using Delly v-0.7.8. *AR* downstream transcriptional activity was determined as AR score. Furthermore, clonal somatic mutations were detected from 25 tumor samples using Mutect2 and Sclust. On the other hand, relationship among metastases in a patient was determined applying SCRATCH algorithm on copy number transition points. Congruence analysis between autosomal- and chromosome X-based SCRATCH relationship of metastases from patient CA34 and CA63 showed that those relationships are not similar by chance.

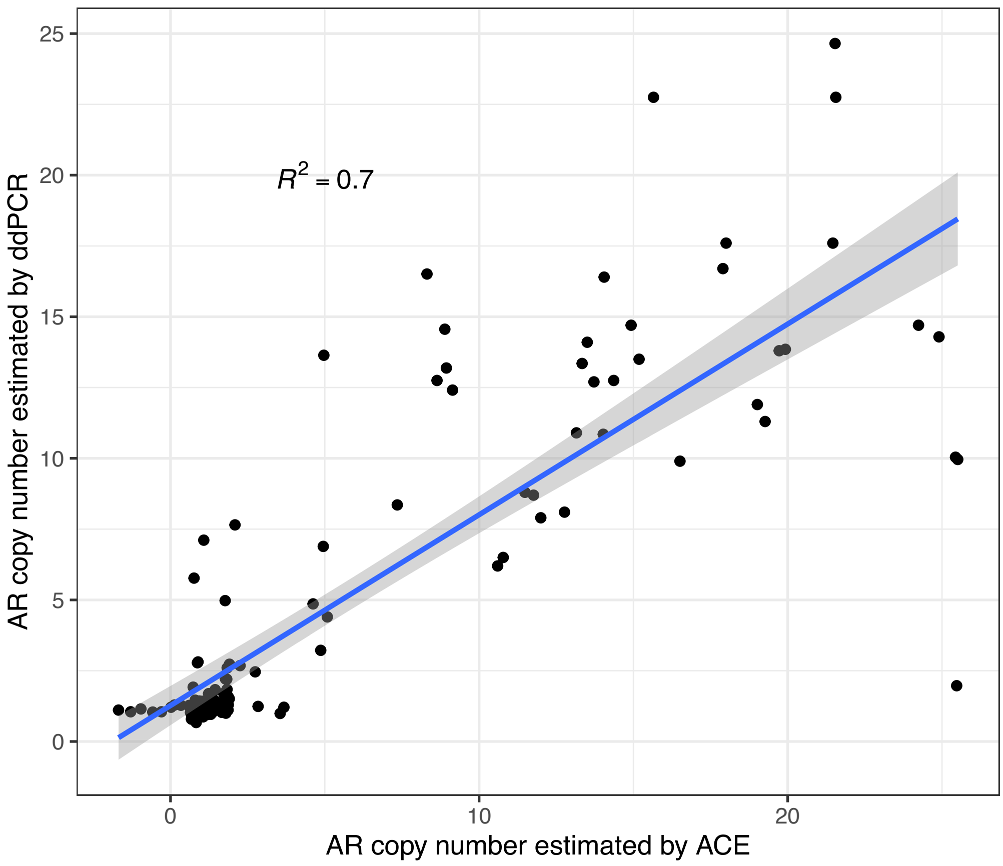


**Figure S2: *AR* copy number estimation**

*AR* copy numbers for 127 samples estimated by droplet digital PCR (ddPCR, along y-axis) correlated highly with the *AR* copy number estimates from shallow whole genome sequencing (using ACE algorithm, along x-axis). Linear regression line is in blue and confidence interval is shown in grey.

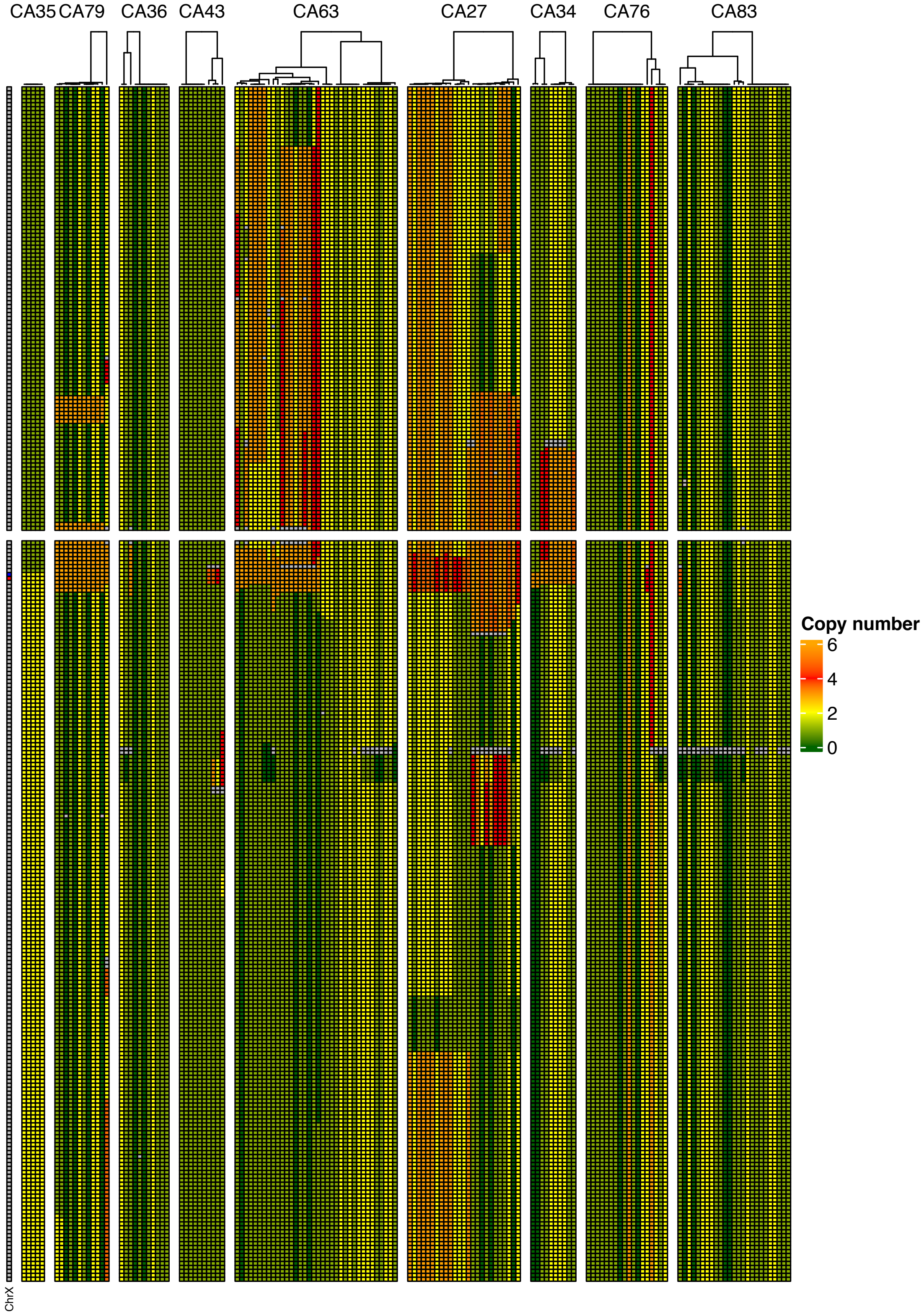
CA35
CA79
CA36
CA43
CA63
CA27
CA34
CA76
CA83
ChrX
Copy number
6
4
2
0

**Figure S3. Copy number gain extends over a large area on chromosome X.**

Heatmap showing the copy number of individual chromosome X bins (500 kb wide) for all 142 samples harvested post-mortem. Color scheme for the copy number shown in the legend. Chromosome X is shown on the left of the heatmap along with its bins covering *AR* gene (in red) and *AR*-associated centromeric enhancer (in blue). Metastases are ordered by patient identification and within patients, based on hierarchical clustering (shown as dendrogram) of correlation distances calculated among copy number transitions on chromosome X.

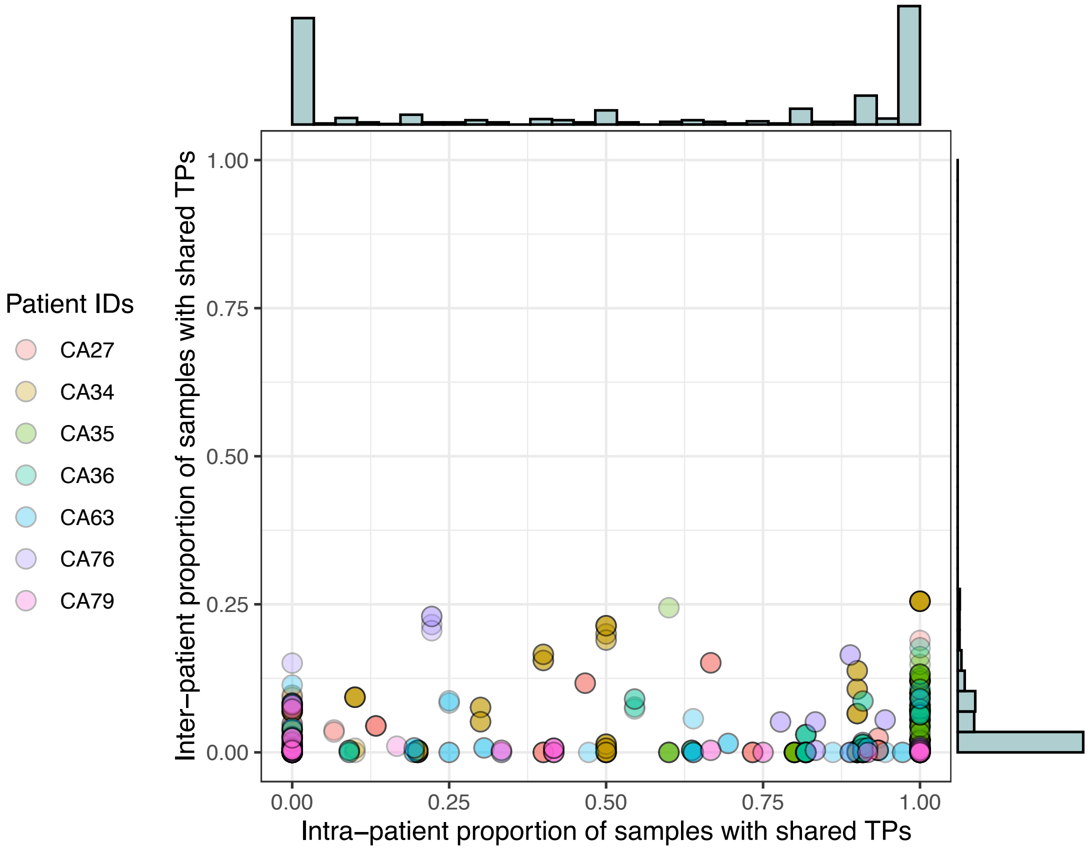


**Figure S4: Overlap of copy number transition points between post-mortem tumor samples and diagnostic biopsies within and across patients**

Circles representing proportion of transition points (TPs) detected in a diagnostic biopsy sample (n=7 patients) that are shared with samples harvested post-mortem from the same patient (x-axis) or any metastases harvested from other patients (y-axis). Marginal histograms confirm that whilst occasional TPs are shared with metastases from other patients, all patients' diagnostic samples share a high proportion of TPs (common or truncal TPs) with metastases from the same patient.

CA34

| Organ | Cluster1 | Cluster2 |
|---|---|---|
| Liver | 4 | 4 |
| Lymph Node | 1 | 1 |
| Plasma | 0 | 1 |

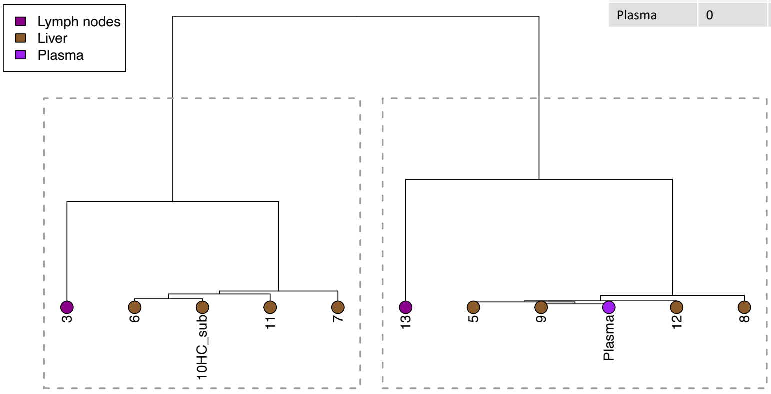


CA36

| Organ | Cluster1 | Cluster2 |
|---|---|---|
| Bone | 0 | 1 |
| Brain | 1 | 0 |
| Dura | 2 | 0 |
| Liver | 6 | 0 |
| Lung | 0 | 1 |
| Plasma | 1 | 0 |

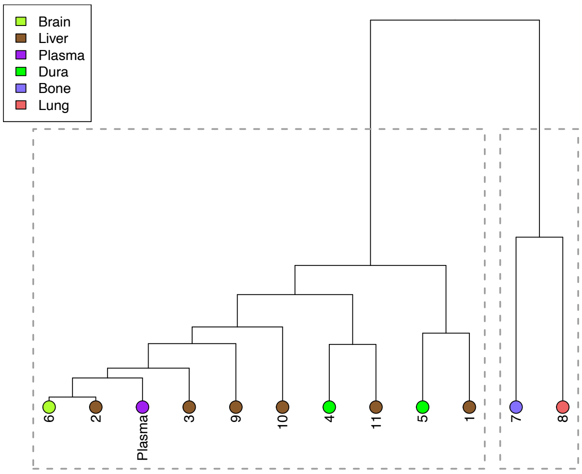

## CA76

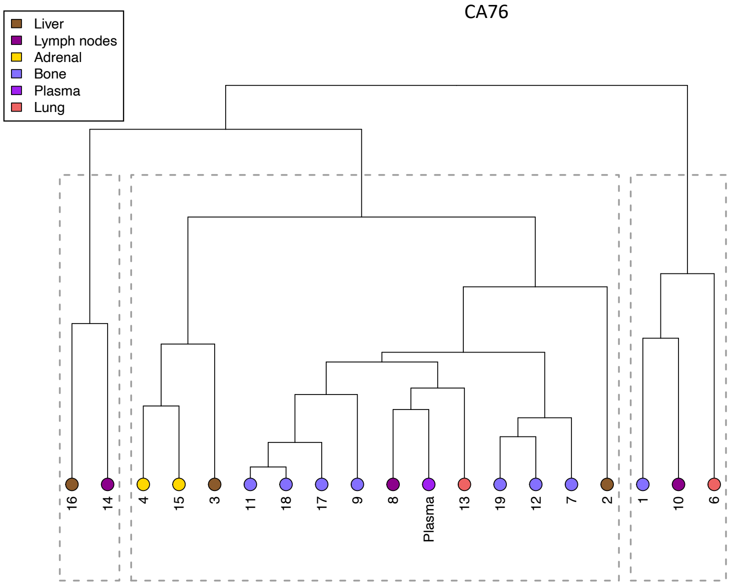


| Organ | Cluster1 | Cluster2 | Cluster3 |
|---|---|---|---|
| Adrenal | 0 | 2 | 0 |
| Bone | 0 | 7 | 1 |
| Liver | 1 | 2 | 0 |
| Lung | 0 | 1 | 1 |
| Lymph Node | 1 | 1 | 1 |
| Plasma | 0 | 1 | 0 |

## CA83

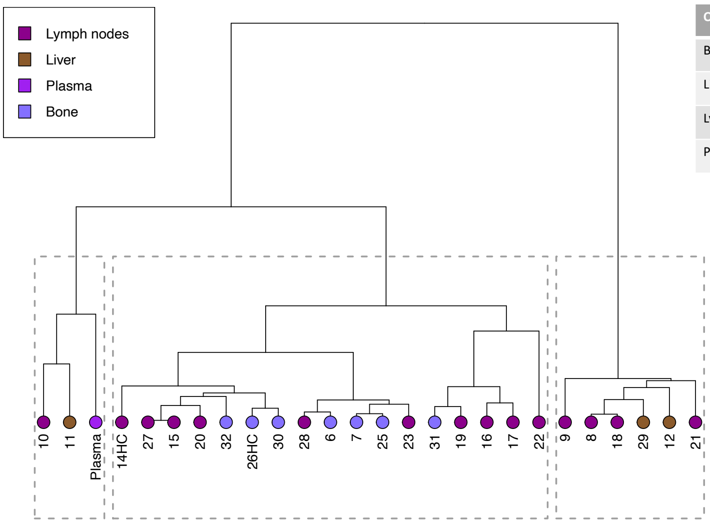


| Organ | Cluster1 | Cluster2 |
|---|---|---|
| Bone | 7 | 0 |
| Liver | 1 | 2 |
| Lymph Node | 11 | 4 |
| Plasma | 1 | 0 |

**Figure S5: SCRATCH relational networks of autopsy tumors for four patients with their plasma collected post-mortem**

SCRATCH relational network (described in methods), built on copy numbers at autosomal transition points, derived from the post-mortem metastatic samples and plasma (tumor content >= 0.2) collected at death from four patients (CA34, CA36, CA76 and CA83), are shown. The circles at the terminal nodes depict organ site for the metastatic samples or plasma in a patient. Metastatic samples and plasma are grouped in two to three clusters (details in methods) and separated by dashed grey boxes. Inset tables show the distribution of organ sites of metastases and plasma in individual clusters.

**CA27**

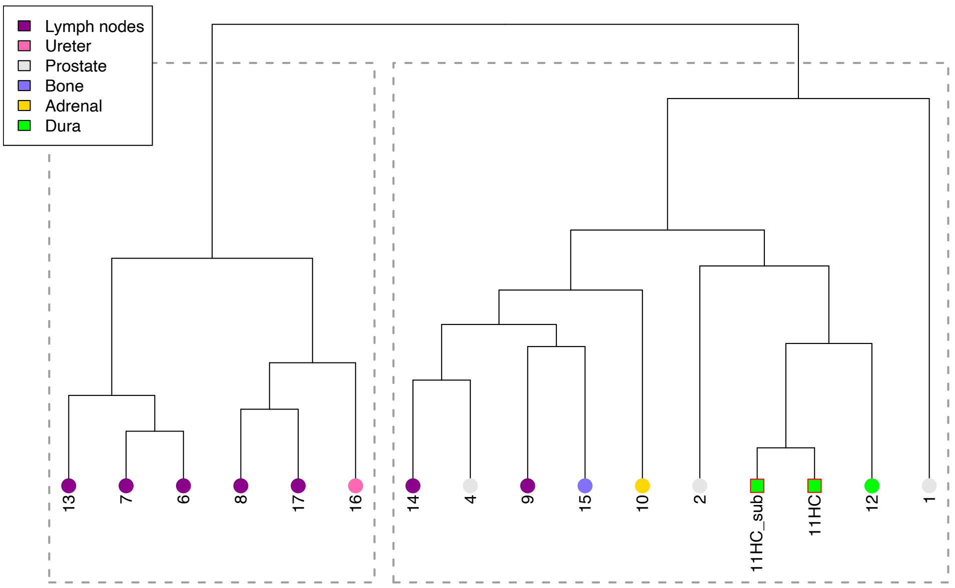
Lymph nodes
Ureter
Prostate
Bone
Adrenal
Dura
13
7
6
8
17
16
14
4
9
15
10
2
11HC_sub
11HC
12
1

**CA34**

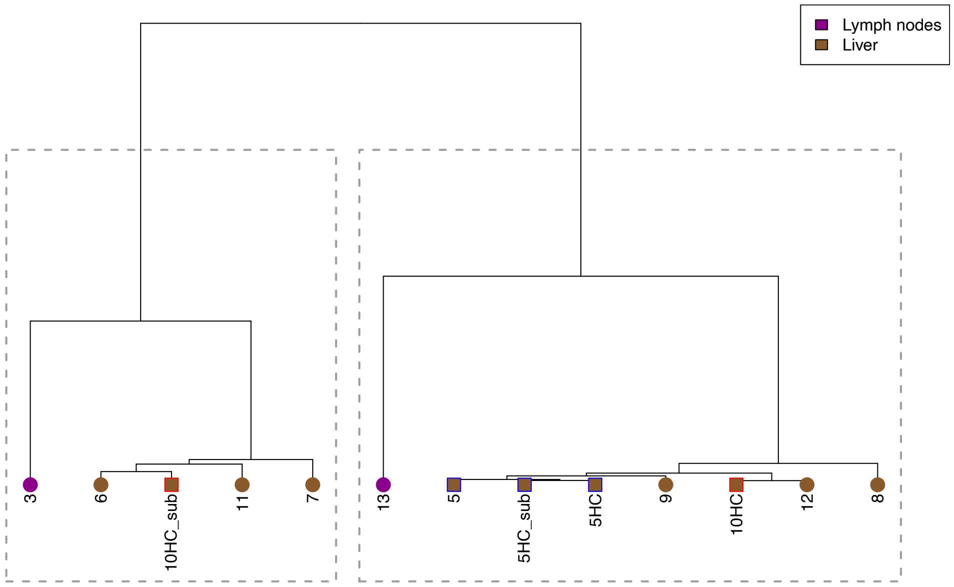
Lymph nodes
Liver
3
6
10HC_sub
11
7
13
5
5HC_sub
5HC
9
10HC
12
8

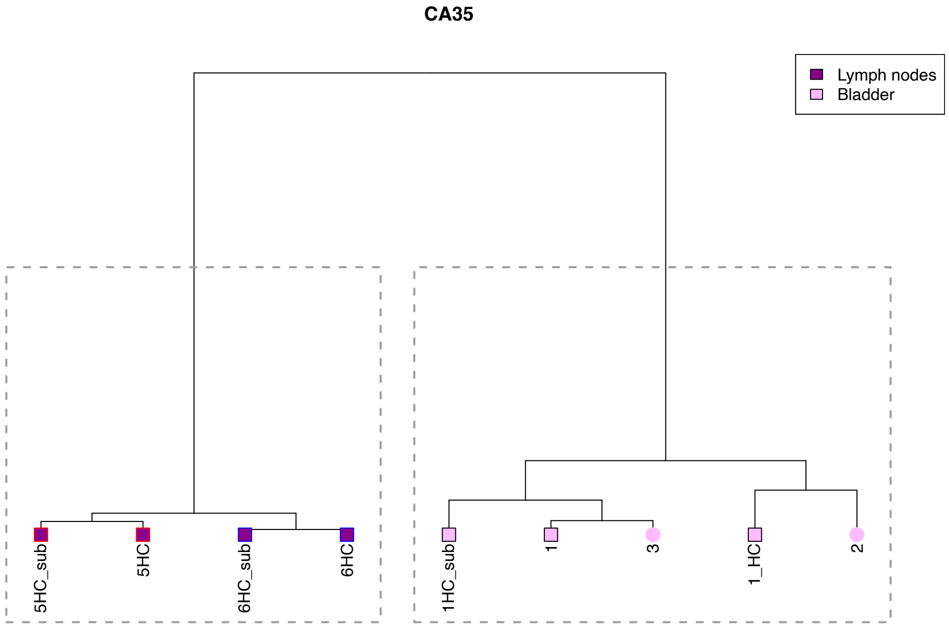
CA35
Lymph nodes
Bladder
5HC_sub
5HC
6HC_sub
6HC
1HC_sub
1
3
1_HC
2

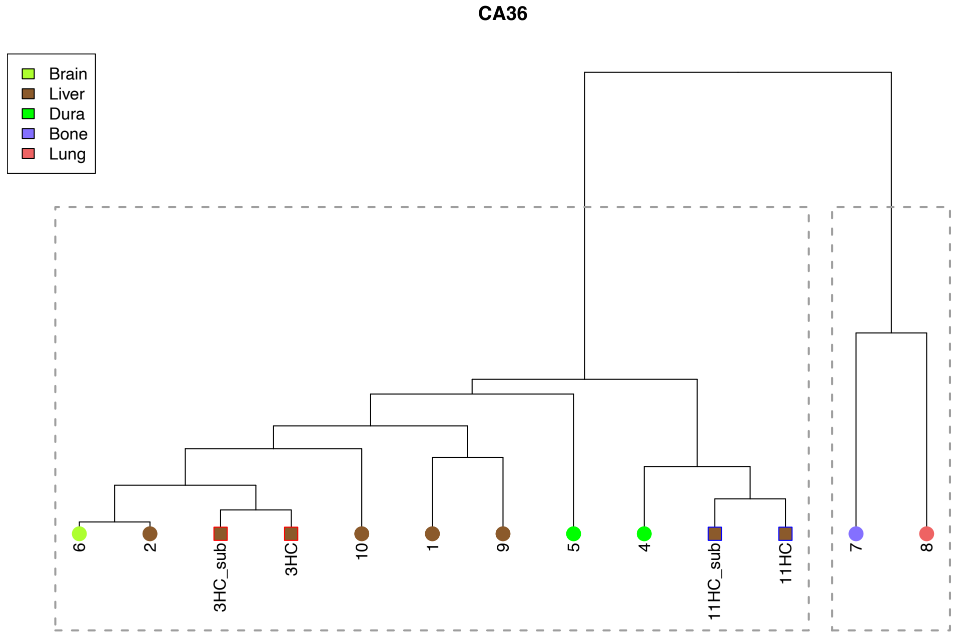
CA36
Brain
Liver
Dura
Bone
Lung
6
2
3HC_sub
3HC
10
1
9
5
4
11HC_sub
11HC
7
8

**CA43**

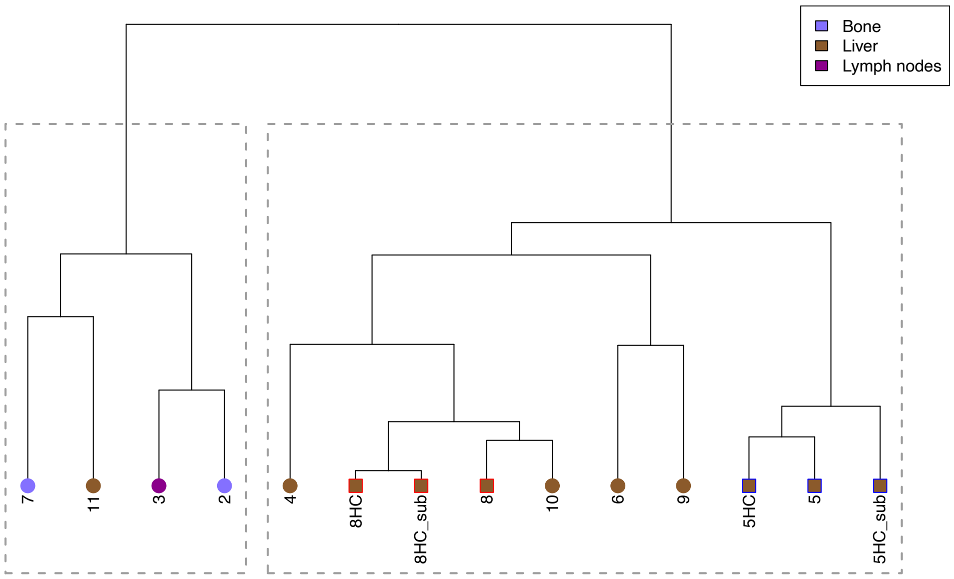
Bone
Liver
Lymph nodes
7
11
3
2
4
8HC
8HC_sub
8
10
6
9
5HC
5
5HC_sub

**CA63**

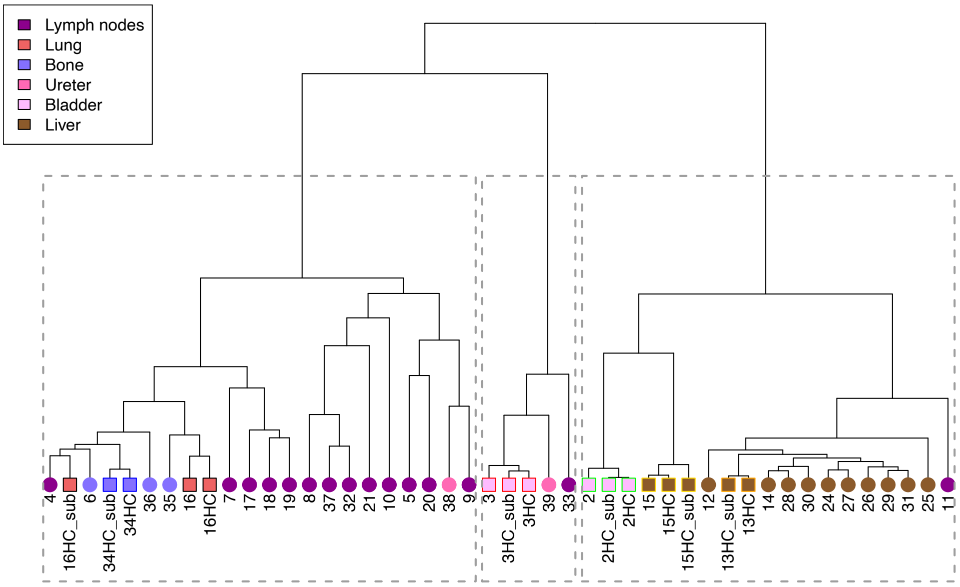
Lymph nodes
Lung
Bone
Ureter
Bladder
Liver
4
16HC_sub
6
34HC_sub
34HC
36
35
16
16HC
7
17
18
19
8
37
32
21
10
5
20
38
9
3
3HC_sub
3HC
39
33
2
2HC_sub
2HC
15
15HC
15HC_sub
12
13HC_sub
13HC
14
28
30
24
27
26
29
31
25
11

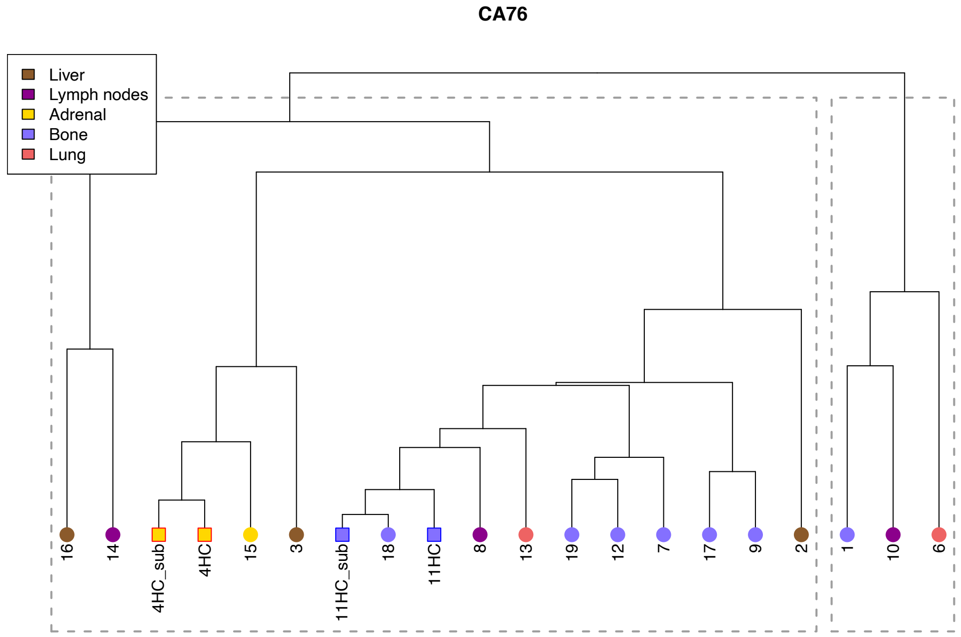
CA76
Liver
Lymph nodes
Adrenal
Bone
Lung
16
14
4HC_sub
4HC
15
3
11HC_sub
18
11HC
8
13
19
12
7
17
9
2
1
10
6

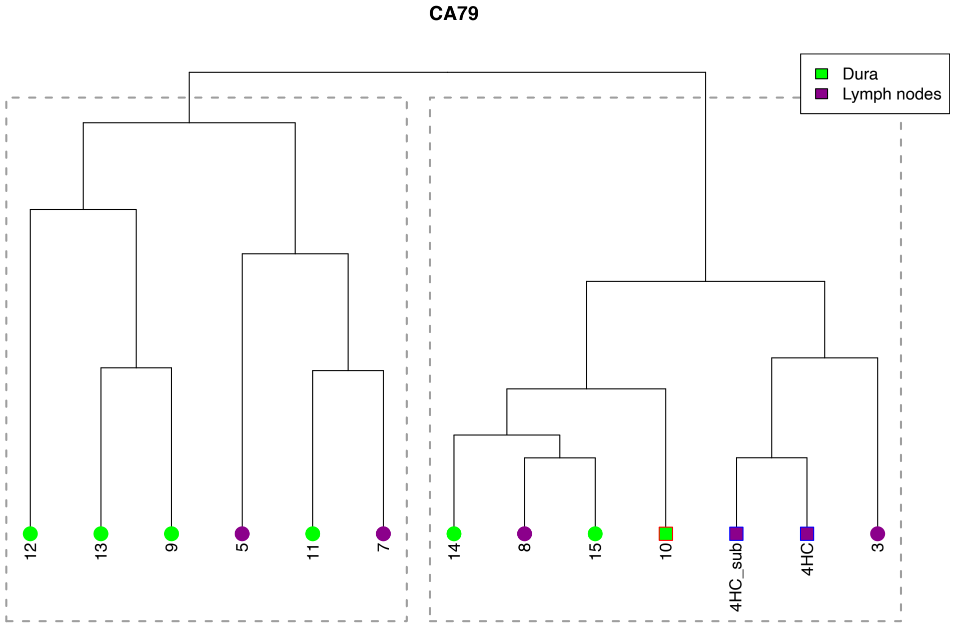
CA79
Dura
Lymph nodes
12
13
9
5
11
7
14
8
15
10
4HC_sub
4HC
3

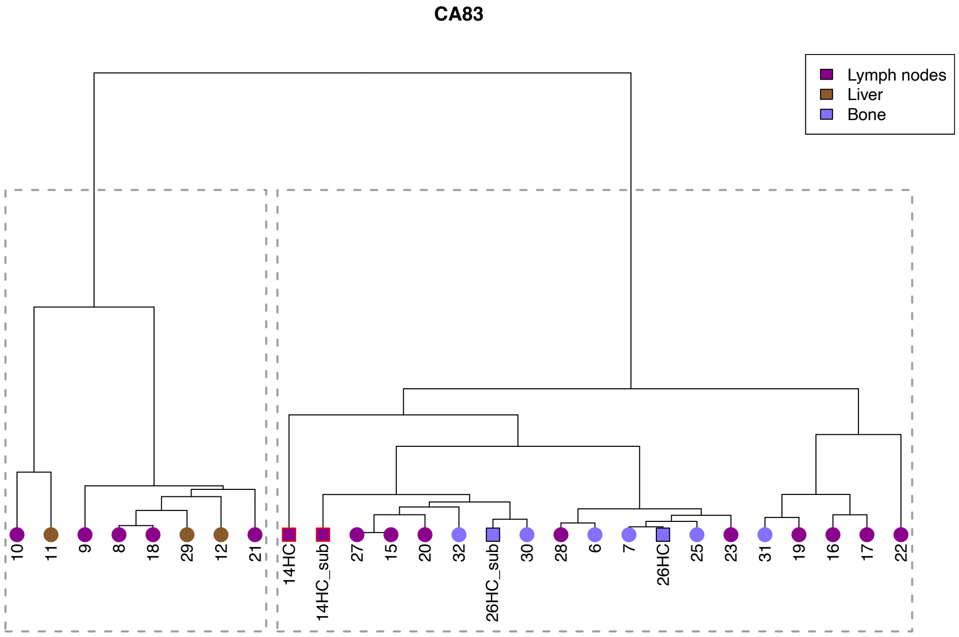


**Figure S6: Relationship assignment of metastases on a SCRATCH relational network is not influenced by sequencing depth**

In SCRATCH relationship networks, tumors with high coverage sequencing (“HC”) and/or subsampled version of them (“HC_sub”) were assigned predominantly close (or in the same cluster, denoted by grey dashed boxes) to the same samples with low coverage sequencing data. The organ sites are depicted with color-coded circles (color scheme in the legend) on the terminal nodes and the samples in comparison (outlined with same-colored lines) are depicted in same-colored squares.

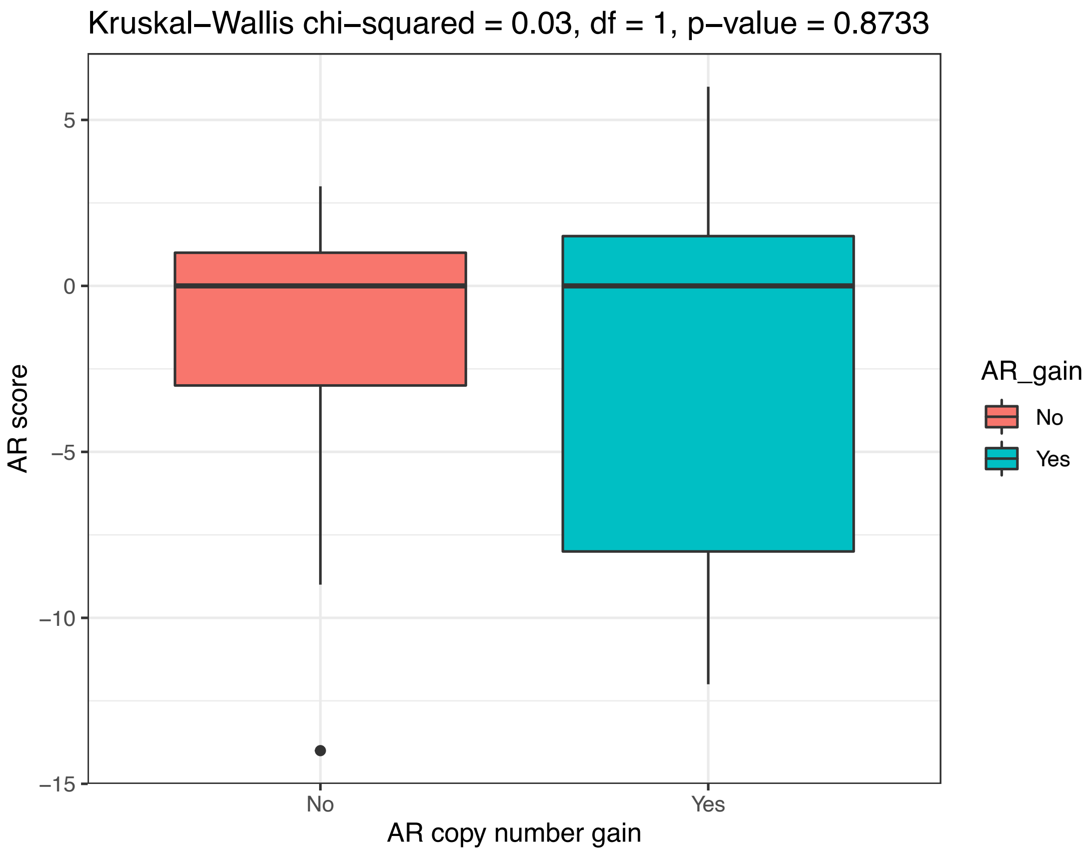


**Figure S7: Expression of AR-target genes are not directly correlated with *AR* expression**

Boxplots of the expression of AR-regulated genes (calculated as AR-score) in metastases with *AR* copy number gain or no gain (significance of difference calculated using Kruskal-Wallis test).
